# Optical characterization of molecular interaction strength in protein condensates

**DOI:** 10.1101/2024.03.19.585750

**Authors:** Timon Beck, Lize-Mari van der Linden, Wade M. Borcherds, Kyoohyun Kim, Raimund Schlüßler, Paul Müller, Titus Franzmann, Conrad Möckel, Ruchi Goswami, Mark Leaver, Tanja Mittag, Simon Alberti, Jochen Guck

## Abstract

Biomolecular condensates have been identified as a ubiquitous means of intracellular organization, exhibiting very diverse material properties. However, techniques to characterize these material properties and their underlying molecular interactions are scarce. Here, we introduce two optical techniques – Brillouin microscopy and quantitative phase imaging (QPI) – to address this scarcity. We establish Brillouin shift and linewidth as measures for average molecular interaction and dissipation strength, respectively, and we used QPI to obtain the protein concentration within the condensates. We monitored the response of condensates formed by FUS and by the low-complexity domain of hnRNPA1 (A1-LCD) to altering temperature and ion concentration. Conditions favoring phase separation increased Brillouin shift, linewidth, and protein concentration. In comparison to solidification by chemical crosslinking, the ion-dependent aging of FUS condensates had a small effect on the molecular interaction strength inside. Finally, we investigated how sequence variations of A1-LCD, that change the driving force for phase separation, alter the physical properties of the respective condensates. Our results provide a new experimental perspective on the material properties of protein condensates. Robust and quantitative experimental approaches such as the presented ones will be crucial for understanding how the physical properties of biological condensates determine their function and dysfunction.

## 1 Introduction

Cells organize their complex inner structure to control the reactions of different components in space and time. This can be achieved by enclosing specific parts of the cellular interior within a membrane, as is the case for the nucleus, mitochondria, or transport vesicles. Another way, which has attracted increasing interest in recent years, is the formation of membraneless compartments via phase separation. Phase separation of macromolecules into a condensed phase occurs above threshold concentrations of the macromolecules, and the driving force for phase separation is modulated by the concentration of ions, molecular crowders, and ligands, and by physical parameters such as temperature [1, 2].

Due to their distinct molecular composition, biomolecular condensates may provide an environment with particular physical properties impacting the chemical and physical processes within the condensates [3, 4]. Some condensates are highly dynamic accumulations of certain proteins and nucleic acids promoting specific biochemical reactions, such as the nucleolus [5] or DNA damage repair sites [6, 7]. However, there is also a continuum of more stable, less dynamic condensates [3], for example, Balbiani bodies during cell dormancy [8], or keratohyalin granules which are crucial for skin barrier formation [9]. Moreover, the condensates’ material properties are variable and may change due to the partitioning of different molecules [10, 11], or in response to physical parameters like temperature [12]. Condensates that are viscoelastic fluids initially, may also transition into viscoelastic solids [12] or solid cross-beta fibrils [6, 13] over time — a state often associated with pathology. For example, mutations in the fused in sarcoma (FUS) and hnRNPA1 proteins investigated in this study are implicated in a spectrum of neurodegenerative diseases, including amyotrophic lateral sclerosis (ALS) [14, 15]. The question of how material properties of biological condensates relate to their diverse functions has been referred to as a major question in the field of biological phase separation [3].

For the characterization of the complex viscoelastic properties that biological condensates may exhibit, or to uncover the underlying molecular interactions, the state of the art comprises the use of a set of techniques reporting on different characteristics. A molecular perspective, that is for example taken by nuclear magnetic resonance (NMR) [16] or Förster resonance energy transfer (FRET) [17], helps to understand single residue dynamics and specific interaction patterns. For a more general characterization of bulk material properties, however, different approaches are needed. A common method used in the field is fluorescence recovery after photobleaching (FRAP). Assuming a purely viscous fluid, FRAP can be used for a qualitative assessment of relative viscosity [18]. Challenges of this technique mainly arise due to the required assumptions about the system probed and the consequential difficulty obtaining robust quantitative values [19]. Condensate coalescence times are typically utilized to determine inverse capillary velocities, which can be converted to surface tensions with the knowledge of viscosities – again assuming a purely viscous fluid [20]. Optical traps [18, 21, 22] or tracking of tracer particles [23] allow the extraction of complex elastic shear moduli, providing direct insight into the viscoelastic properties of condensates. The incorporation of beads or tracer particles into the condensates, however, can be experimentally cumbersome and potentially influence the results obtained.

Moreover, a simple measure of molecular interaction strengths applicable to all states of matter found in biological condensates is not yet available. With these challenges in mind, we chose a purely optical approach that does not rely on fluorescent labels or extensive assumptions and reports on macroscopic material properties and average molecular interaction strength. We employed Brillouin microscopy to investigate biological phase separation at intermediate length and timescales by quantifying the average molecular interaction strength that drives collective molecular motion. One form of collective molecular motion is a thermally excited pressure wave (Figure 1) propagating through a material at the speed of sound, *v*. Light that is scattered by such a propagating wave experiences a frequency shift ν_B_, called Brillouin shift. This type of inelastic light scattering was described theoretically by Brillouin and Mandelstam [24, 25] in the 1920s. The absolute value of the Brillouin shift is proportional to the speed of sound (ν_B_ ∝ *v*), which is determined by longitudinal modulus *M* and mass density *ρ* as 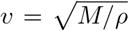. The longitudinal modulus can be interpreted as an average molecular coupling constant, giving rise to a resistance against uniaxial compression. Important to note is that this resistance is not specific to a particular type of molecule (e.g., proteins); any molecule within the probed volume contributes. A stronger coupling between molecules gives rise to a higher sound velocity and a greater Brillouin shift 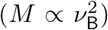 The molecular motion probed takes place in the sub-nanosecond regime and averages molecular interactions over a few hundred nanometers.

**Figure 1:**
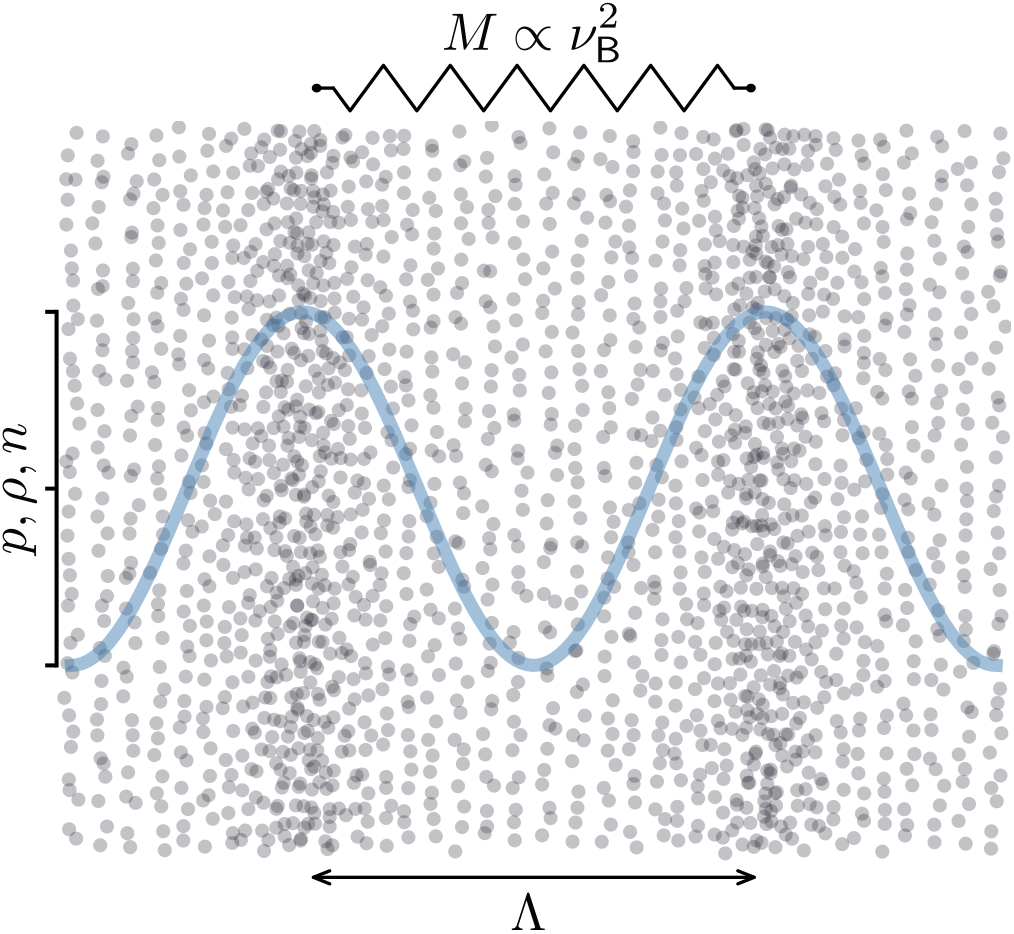
Illustration of a thermally excited pressure wave. Individual particles are indicated as grey dots, the blue line indicates the change in pressure, density, and refractive index compared to an unperturbed state. The amplitude of the wave and the corresponding particle displacement are exaggerated for better visualization. Due to the pressure wave, particles get pushed together (high pressure) or pulled apart (low pressure). (Inter)molecular interactions give rise to an inherent resistance to this collective particle displacement. This resistance can be quantified by the longitudinal modulus *M*, an average molecular coupling constant on the length scale of the acoustic wavelength Λ, which is in our case on the order of a few hundred nanometers corresponding to an acoustic frequency in the Giga-Hertz regime. Stronger particle interactions lead to a higher longitudinal modulus and faster pressure wave propagation. As the pressure *p* is coupled to density *ρ* and consequently refractive index *n*, periodic changes in pressure can scatter light (Brillouin scattering). The measurement of the Brillouin shift ν_B_ gives access to the longitudinal modulus and therefore the average molecular interaction strength in the probed volume. The wave attenuation, on the other hand, is determined by the dissipative properties of the medium and is linked to the Brillouin linewidth Δ_B_. The relation of these quantities to other established viscoelastic quantities is further treated in the supplement (S 5).

While the Brillouin shift is associated with the elastic properties of the material (i.e., the longitudinal modulus), the linewidth Δ_B_ is related to dissipation. In a liquid that is homogeneous on the length scale of the sound wave, sound attenuation is determined by the internal degrees of freedom of the material that dissipate energy and give rise to its viscosity (see also Section S 5) [26, 27].

While the application of Brillouin microscopy for biophysics research is a relatively recent development [26, 28], the investigation of viscoelasticity with sound waves has a long history. Ultrasonic investigations were used broadly in material science and polymer physics, as well as to study phase separation [29, 30]. One specific advantage of using a sound-based method to study protein condensates is the possibility to quantitatively characterize material properties in any state of matter. There is no limitation to liquids or solids; in the case of Brillouin microscopy, optical accessibility is the only requirement. Furthermore, the determination of mechanical properties does not involve the application of a mechanical model which requires certain assumptions about the system probed. This is particularly advantageous for samples where the state of matter is not known or changes over time.

As the calculation of the longitudinal modulus based on the Brillouin shift requires knowledge about the refractive index (see 4.5), a combination of Brillouin microscopy and quantitative phase microscopy such as QPI or optical diffraction tomography (ODT) is advantageous [31]. Those techniques are able to capture the complex optical fields, including the phase information, while conventional light microscopes only probe the light intensity. The phase information can be converted into the sample’s refractive index distribution, if the object geometry is known (e.g. spherical), or if the specimen is imaged from different angles in a tomographic manner [32, 33]. Furthermore, the refractive index can be utilized to calculate protein concentration, volume fraction, as well as mass density, which is also required for longitudinal modulus computation (see 4.5).

Brillouin microscopy has already been applied to membraneless organelles. While other groups have previously studied nucleoli [34] and FUS-condensates in fixed cells [35], our group has recently employed a combination of spatially resolved refractive index measurements and Brillouin microscopy to study nucleoli, FUS-condensates, and poly-Q accumulations in living cells. A key finding was that nucleoli and poly-Q aggregates have a lower compressibility and higher refractive index than the surrounding material [31]. Following this proof of principle showing the applicability of Brillouin microscopy for the investigation of biomolecular condensates in general, a further establishment of this experimental method needed additional experimental work evaluating the sensitivity of this technique to changes in the physical state of the condensates in a systematic manner. Therefore, we tuned parameters known to affect phase separation and quantified their impact on the physical properties of different protein condensates.

Here, we used a combination of Brillouin microscopy and quantitative phase imaging (QPI) to investigate the impact of solution conditions, temperature, time, and amino acid sequence alterations on protein condensates in vitro, in a non-invasive, purely optical manner. We quantified changes in the molecular interaction strength in terms of Brillouin frequency shift, and dissipative properties in terms of Brillouin linewidth. Additionally, we compute protein concentration and volume fraction from refractive index measurements. We found thatprotein condensates exhibited higher values in the aforementioned optomechanical quantities in conditions with a stronger tendency for phase separation (i.e., low temperatures [36, 37], high, or low salt concentrations [38]) revealing stronger molecular interactions. We also characterized the aging process of FUS condensates and found that their protein concentration and volume fraction, as well as their dissipative properties, remained unaffected. The aging process was also characterized by a gradual increase in Brillouin shift (i.e., molecular interaction strength), however, this effect was much weaker than the Brillouin shift elevation due to chemical fixation. Accordingly, the observed aging process may be characterized by a progressive establishment of a crosslinked polymer network across the condensates. Finally, we showed that the molecular grammar governing macromolecular condensation also has a strong impact on the physical properties of the condensates. Our experiments with variants of the low-complexity domain of the hnRNPA1 protein (A1-LCD) [39] revealed distinct molecular interaction and dissipation strength, as well as different densities. Moreover, we found that the driving force for phase separation (quantified by the saturation concentration) showed a correlation with the molecular interaction strength in the condensates.

Overall, we demonstrated an all-optical approach that does not rely on fluorescent labels to quantify molecular interaction strength as well as protein concentration of protein condensates. We showed the sensitivity of our measurement techniques to known parameters affecting phase separation and physical properties of protein condensates. Such quantitative approaches are crucial to consolidate the current state of knowledge and to address open questions in the field of biological phase separation.

## 2 Results

### 2.1 Experimental setup

Figure 2 illustrates the experimental setup as well as the analysis workflow. Brillouin microscopy was performed by doing point measurements in the condensed and dilute phases of phase-separated protein solutions. Surface wetting of the protein was minimized by polyethylene glycol coating, keeping the condensates in a spherical shape. This enabled the use of QPI to measure the refractive index and calculate protein concentration and volume fraction. Furthermore, the refractive index was utilized to calculate mass density and longitudinal modulus (see Sections 4.4 and 4.5).

**Figure 2:**
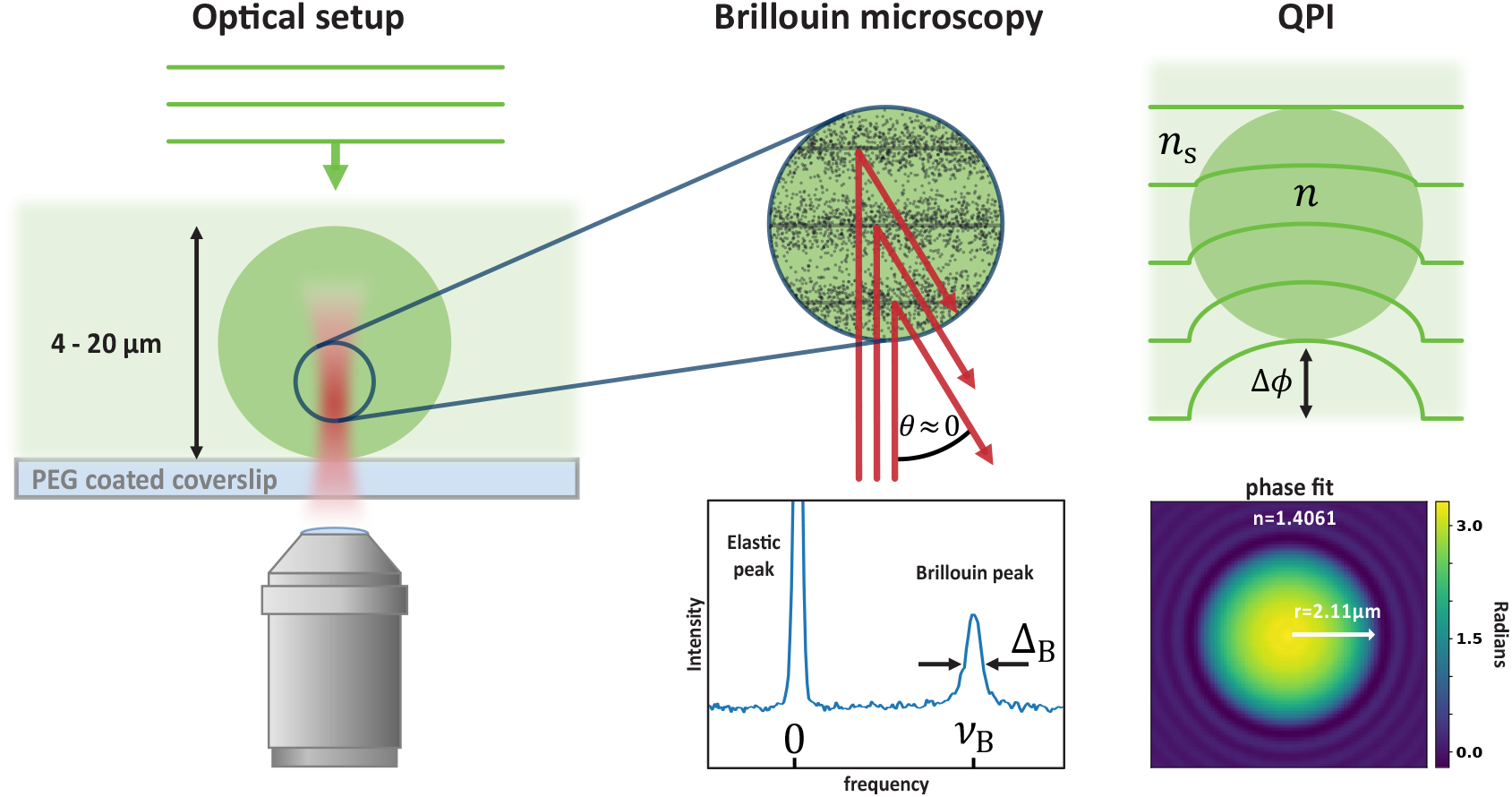
Introduction of experimental techniques. Left column: Schematic representation of the optical setup. Brillouin microscopy probes a single confocal spot, QPI illuminates the complete field of view with a plane wave. Middle column: Schematic representation of the Brillouin scattering process in the confocal volume probed, where *θ* denotes the scattering angle. The incoming probe beam is inelastically scattered by microscopic pressure waves. Brillouin frequency shift ν_B_ (i.e., the distance between the elastic peak and the Brillouin peak on the frequency axis) and linewidth Δ_B_ are indicated in an example spectrum of water. Right column: QPI quantifies the phase delay Δ*ϕ* introduced by a specimen compared to the surrounding medium with a known refractive index *n*_s_. Due to the higher refractive index of the specimen *n > n*_s_, the light traveling through the specimen is delayed resulting in a distortion of the incoming plane wave. This distortion can be captured in a phase image showing Δ*ϕ*. A custom algorithm fits the phase image obtained, assuming a spherical shape of the specimen, providing the radius *r* and refractive index *n* of the fitted sphere (see Section 4.3 for more details). Non-spherical samples require a tomographic imaging techique such as optical diffraction tomography (ODT).

### 2.2 Increasing temperature weakens molecular interaction strength in a polymer rich-phase

Temperature is an important thermodynamic control parameter for the formation of protein condensates. Previous reports have revealed that FUS condensation follows an upper critical solution temperature (UCST) phase diagram [37]. Therefore, we asked how temperature increase, which diminishes the driving force for phase separation [36, 37], affects the physical properties of FUS condensates.

To obtain a reference system illustrating the impact of temperature and solute concentration on the average molecular interaction and dissipation strength of a polymer solution, we first measured the Brillouin shift and linewidth of polyethylene glycol 3350 (PEG) solutions. We found that increasing the temperature of water (0% PEG) or a 10% PEG solution increased Brillouin shifts while decreasing linewidths (Figure 3A). Above a PEG concentration of 20%, the slope Δ_B_ vs. ν_B_ became positive, yielding both an increased Brillouin shift and linewidth with lower temperatures. Our results are in line with previous Brillouin scattering measurements of PEG 600 [40] and concentrated sucrose solutions [41].

**Figure 3:**
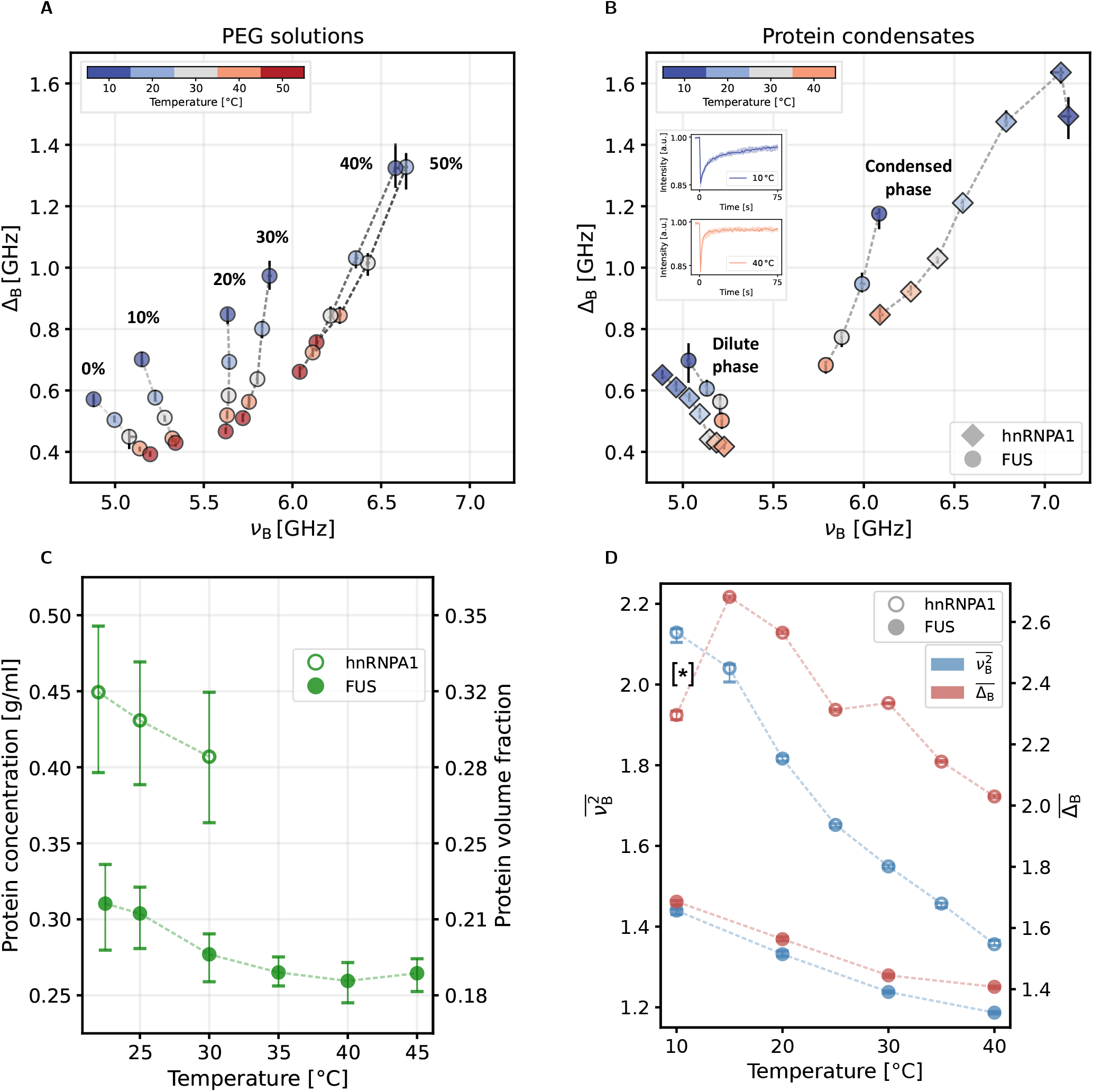
Temperature alterations of PEG solutions and protein condensates. A: Brillouin shift ν_B_ and linewidth Δ_B_ for PEG solutions at different temperatures. The labels indicate the PEG weight fraction. B: Brillouin shift ν_B_ and linewidth Δ_B_ for FUS and A1-LCD condensates and corresponding dilute phase for different temperatures at 150 mM KCl with 5% dextran and 150 mM NaCl, respectively. A control experiment for FUS condensates without the addition of dextran can be found in the supplement (Figure S 8A). Inset shows fluorescence intensity recovery after photobleaching (FRAP) of FUS condensates at 10 °C and 40 °C. The solid line shows the median intensity recovery of five individual condensates. C: Corresponding protein concentration and protein volume fraction of protein condensates for various temperatures. The accessible temperature range for A1-LCD was limited by the onset of condensate dissolution. D: Brillouin shift squared 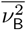 (left axis) and Brillouin linewidth 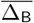 (right axis) of protein condensates normalized to the respective values of the dilute phase. Note that the Brillouin shift of A1-LCD at 10 °C, indicated by [*], might be subjected to a systematic underestimation; more details can be found in Section S 4. Markers indicate median values; error bars represent the range containing 68.3% of the data points that are closest to the median. In some cases, the marker diameter is larger than the error bars.

The increase of the Brillouin shift in a dilute polymer solution due to an elevation of temperature from 10 °C to 50 °C can be explained by the viscoelastic properties of water. Water exhibits a minimum in compressibility (resulting in a maximum of the bulk modulus *K*) at 45 °C [42] and a vanishing shear modulus *G* = 0. Due to the relation 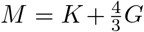, this implies a maximum of the longitudinal modulus [43] and accordingly in the Brillouin shift (see also Section S 5). In summary, the viscoelastic properties of water dominate at low solute concentrations up to 10% PEG, while the solute dominates at concentrations above 20%.

After the establishment of the PEG-model system, we conducted *in vitro* experiments on FUS and A1-LCD condensates. To extend the accessible temperature range we added 5% dextran to the FUS buffer system preventing the dissolution of the condensates. In the dilute phase of the phase-separated protein solution (Figure 3B) we found an increase of the Brillouin shift and a reduction of the linewidth as a result of a temperature increase. The same trend was observed before for water and a 10% PEG solution (Figure 3A). The protein condensates on the other hand (Figure 3B), exhibited a reduction of Brillouin shift upon the same temperature change, similar to a 30% PEG solution (Figure 3A). Figure 3 also illustrates that the impact of polymer concentration on linewidth and Brillouin shift decreases for higher temperatures. This effect was observed for the phase-separated protein solution as well as the PEG model system. To compare our results with a widely used technique in the field, we performed fluorescence recovery after photobleaching (FRAP) to qualitatively inspect protein diffusion in the FUS condensates. We found a faster intensity recovery at higher temperatures (Figure 3B). This observation together with the measured trends in the Brillouin linewidth shows a reduction of dissipation upon temperature increase.

In addition, we investigated whether an increase in temperature would also result in a decrease in refractive index [44] and density [45, 46] as previously reported for PEG solutions. Such behavior would be expected for a narrowing two-phase regime at increasing temperature in a UCST system. Indeed, we observed a subtle decrease in refractive index and protein concentration of FUS condensates upon a temperature increase from 25 °C to 35 °C (Figure 3C), similar to what was reported for mEGFP-TAF15 condensates [47]. The same trend was found for A1-LCD, but the accessible temperature range was more restricted, as we did not use a crowding agent in this case. The onset of condensate dissolution above 30 °C and the consequent lack of spherical condensates prevented QPI measurements at higher temperatures.

As the approximation *ρ/n*^2^ ≈ *const*. is justified in our case (see Section S 6 for further details), it follows that 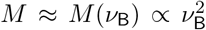 and a measurement of the Brillouin shift is sufficient to detect changes in the longitudinal modulus. Accordingly, we report 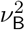 instead of *M* as a measure of molecular interaction strength in the following text. To facilitate the comparison of the different factors that were investigated in this study (temperature, ion concentration, and aging), we utilized the dilute phase for the normalization of Brillouin shift squared 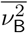 and linewidth 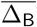 of the dense phase. This approach helps to highlight the distinct changes of the physical properties of the dense phase compared to the dilute phase. Similar normalized quantities employing water as a reference were earlier introduced as “Brillouin elastic contrast” for the Brillouin shift and as “Brillouin viscous contrast” for Brillouin linewidth, respectively [48].

Plotting these normalized quantities for the phase-separated FUS system as a function of temperature shows a clear reduction of 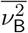 and 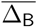 upon temperature increase (Figure 3D). The local maximum of Brillouin linewidth for A1-LCD condensates at 15 °C together with the monotonic trend in the Brillouin shift could denote a relaxation process [49]; i.e. the timescale of molecular motion was slowed down into the regime of the experimentally accessed timescale of few hundred picoseconds.

In summary, we found a reduction in molecular interaction and dissipation strengths in the protein condensates upon temperature increase, accompanied by a reduction in protein concentration, where there was also evidence for a molecular relaxation process in the A1-LCD condensates indicated by a distinct signature in Brillouin shift and linewidth.

### 2.3 Variation of ion concentration reveals distinct interaction regimes

Another important parameter that can tune the interactions between proteins is the ion concentration in the surrounding medium. Ions have specific effects on protein-protein interactions as they directly affect the various possible interaction mechanisms of amino acids [2] and the solubilities of proteins. A previous study demonstrated that FUS exhibits a reentrant phase behavior, resulting in two phase separating regimes, one at low and another one at high ion concentrations [38]. Also, the phase separation behavior of A1-LCD in response to different ion concentrations has already been investigated. In contrast to the FUS protein, a larger ion concentration leads to a lower saturation concentration of A1-LCD, whereas a reentrant phase separation was not observed [50]. We hypothesized that these ion-dependent interaction regimes should be also reflected in the physical properties of the protein condensates. As above for exploring the effect of altering temperature, we added 5% dextran ensuring the presence of FUS condensates of suitable size across the full range of KCl concentrations.

Figure 4 illustrates the distinct impact of ion concentration on the physical properties of the protein condensates. While the buffer solutions showed a linear dependence of refractive index on ion concentration (see supplement Section S 2 and Figures S 6A and S 6B), as well as the Brillouin shift of the dilute phases (Figures S 6C and S 6D), the dense phases exhibited more complex dependencies. The calculation of protein concentration and volume fraction (Figure 4A), which is based on the refractive index difference of buffer and sample (see Section 4.4), showed that the maximum of protein concentration and volume fraction of FUS protein condensates is found at the lowest KCl concentrations. Towards higher KCl concentrations, in the region where FUS condensates would dissolve without the addition of a crowding agent (i.e., 0.5M and 1M KCl), protein concentration and volume fraction showed a minimum, before slightly increasing again at even higher KCl concentrations. A1-LCD condensates, on the other hand, were found to be generally denser than FUS condensates and showed a further increase of protein concentration and volume fraction upon higher NaCl concentrations.

**Figure 4:**
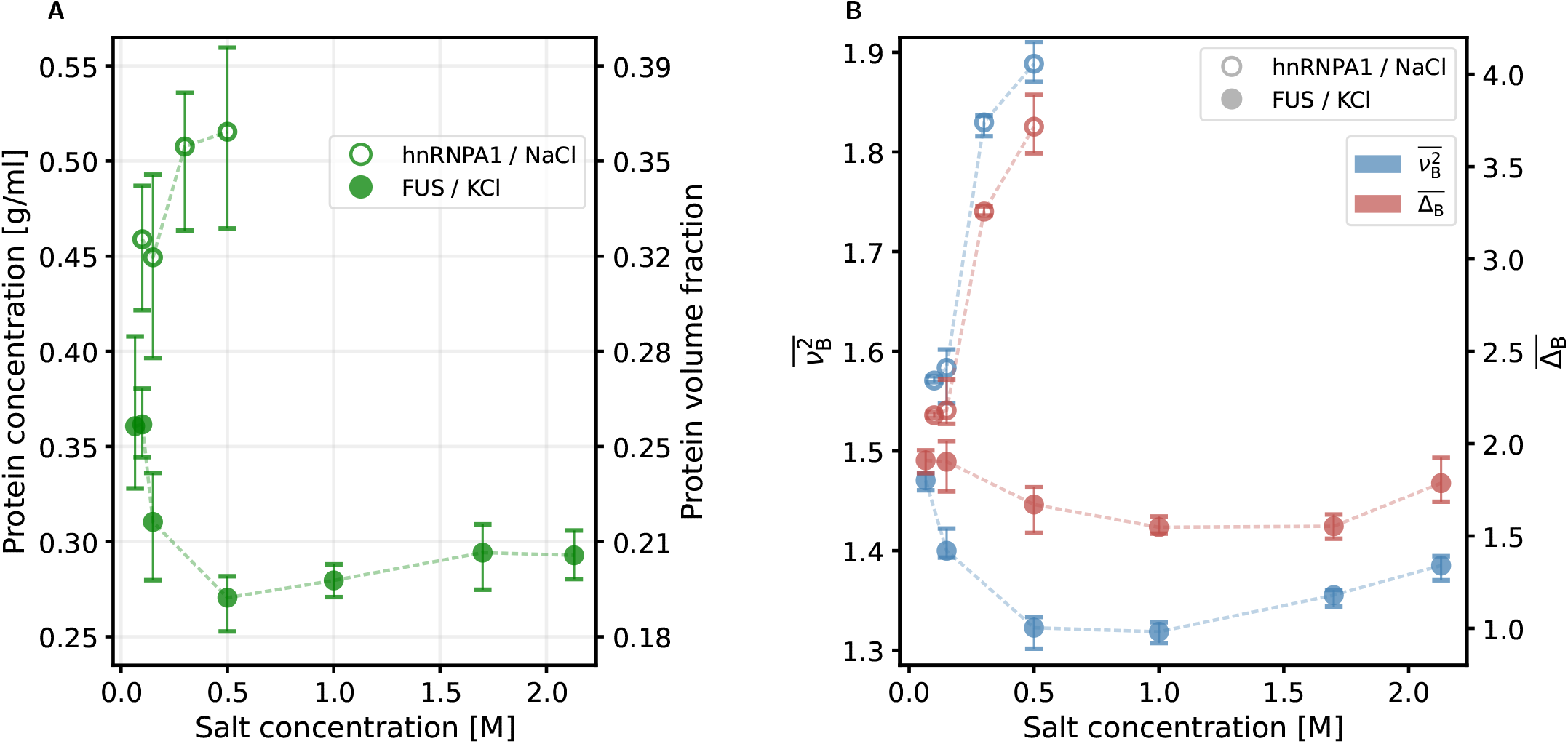
Impact of different ion concentrations on protein condensates. A: Protein concentration and volume fraction of FUS condensates at 5% dextran and different KCl concentrations and of A1-LCD at varying NaCl concentrations. Note that the data shown for FUS at 150 mM KCl is the same as in Figure C at 22.5 °C. 4B: Corresponding Brillouin shift squared 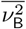 (left axis) and Brillouin linewidth 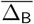 (right axis) normalized to the respective values of the dilute phase. Markers indicate median values; error bars represent the range containing 68.3% of the data points that are closest to the median.

Finally, as for the different temperatures in Figure 3, we computed normalized values for Brillouin shift squared 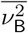 and linewidth 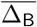 (Figure 4B). For FUS condensates, we identified a minimum in both quantities around a KCl concentration of 1 M, while the largest values were located at the lowest ion concentrations, similar to the trends of protein concentration and volume fraction. In the same line, A1-LCD showed larger values of Brillouin shift and linewidth at higher NaCl concentrations.

In summary, our results show that the physical properties of the condensates change according to the previously identified interaction regimes. In addition, we found that low ion concentrations lead to denser FUS condensates with stronger molecular interactions compared to high ion concentrations.

### 2.4 FUS condensate aging is ion concentration dependent

Multiple publications have investigated the time dependence of viscoelastic properties of FUS protein condensates. In general, the aging process of FUS condensates resulted in reduced protein mobility. The states of matter described for aged FUS condensates range from viscous fluids [51], and mixed liquid-gel states [52], to fibrillar solids [6, 52]. Given this broad range of results, we used Brillouin microscopy and QPI in various buffer conditions to provide an unbiased view of the aging of FUS condensates.

Previous publications reported morphological and mechanical changes in a time frame of about 12 h to 24 h [6, 51]. Accordingly, we conducted QPI measurements shortly after sample preparation, as well as 12 h, 24 h, and 36 h later. In contrast to some previous publications, we could not find morphological changes even after 36 h, other than Ostwald ripening. Accordingly, bigger condensates were growing while smaller ones were shrinking (Figure 5A). In contrast to a previous study on PGL3 condensates reporting an increase in protein concentration over time [47], we could not detect a change in refractive index, protein concentration, or protein volume fraction (Figure 5B).

**Figure 5:**
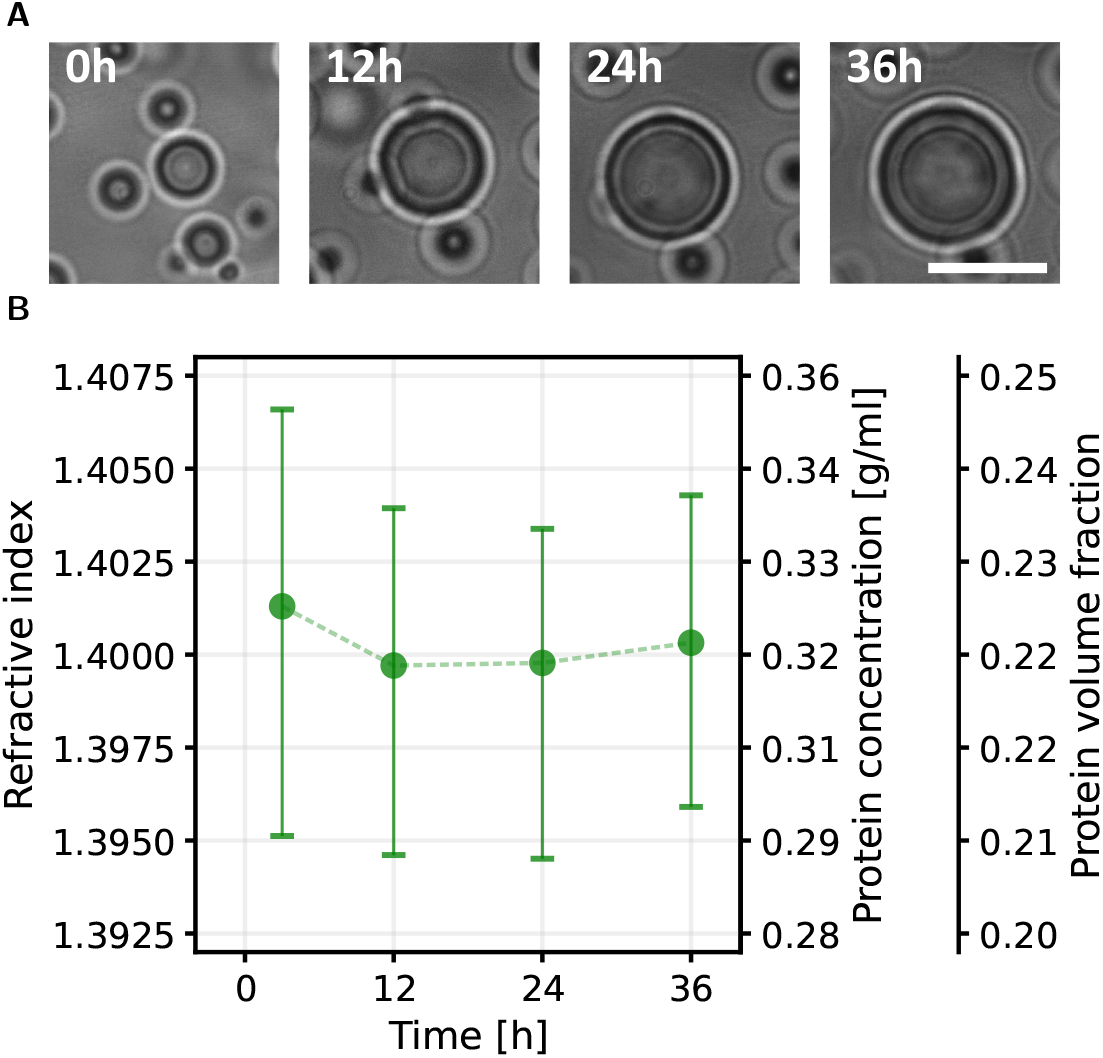
Effect of FUS condensate aging on morphology and density. A: Bright-field images of a representative FUS condensate over a time course of 36 h at 150 mM KCl and 5% dextran. B: Refractive index, protein concentration and volume fraction of FUS condensates measured 3 h, 12 h, 24 h and 36 h after condensate formation. Markers indicate median values. Scale bar: 10 µm.

We also conducted Brillouin microscopy measurements over the same time course as the QPI measurements. We were aiming to investigate the same condensate population over the whole time course. In the case that a particular condensate became inaccessible we chose a new one that was then monitored over the remaining time. To minimize any possible effects of the laser light, we chose a laser power of about 11 mW, even though much higher laser powers (*>* 40 mW at 660 nm) have already been shown to be safe for imaging live cells [53]. As data acquisition was easier for condensates with a diameter of at least 5 µm, due to reflections caused by the refractive index jump at the condensate interface, our measurements were performed mainly on condensates that were growing (Figure 5A).

We found that the aging process of the condensates was characterized by a shallow increase in 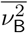 of the condensate population in the case of low ion concentrations. At a KCl concentration of 2.5 M, however, this trend was suppressed (Figure 6A). Similar to the density of the condensates, the dissipative properties of the condensates were also mostly unaffected by the condensate aging (Figure 6B). An increase in viscosity that was reported previously [51] should have been reflected in an increase in Brillouin linewidth.

**Figure 6:**
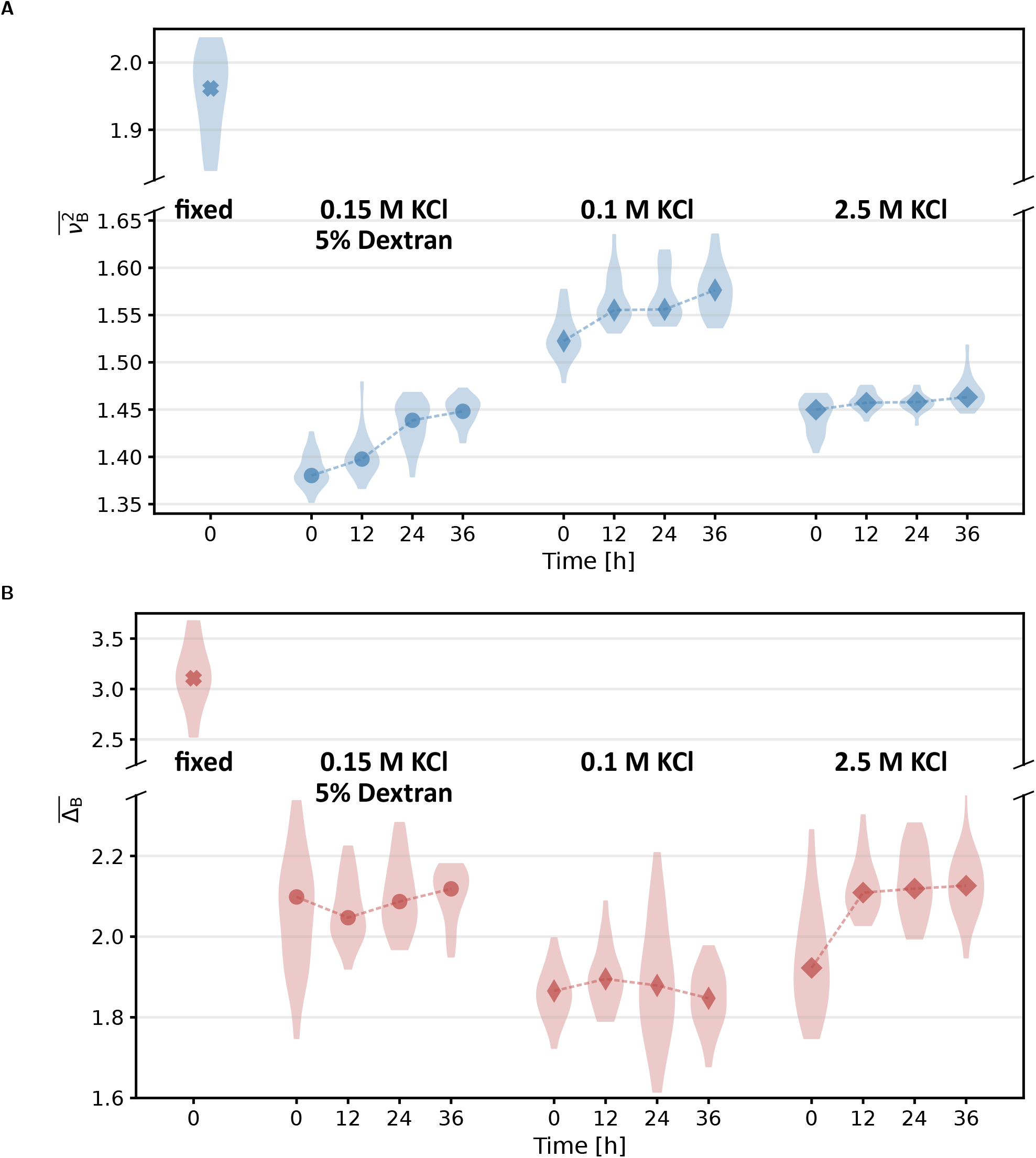
Effect different buffers on FUS droplet aging. Time evolution of 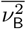 (A) and 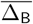 (B) of FUS condensates for three different buffer conditions. Fixed condensates were chemically crosslinked by the addition of 0.05% glutaraldehyde at a KCl concentration of 100 mM. Markers indicate median values.

To create a reference system for a completely solidified protein condensate we added 0.05% glutaraldehyde to a phase separated FUS-protein system. To verify the solidification of the condensates we increased the temperature up to 40 °C after the measurement. As the condensates did not dissolve, we concluded that the solidification was irreversible. The chemical cross-linking led to an increase in 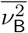 of 29% leading to the highest value of all conditions investigated in this study.

These findings suggest that the observed ion-dependent condensate aging process is not characterized by the formation of ordered crystalline structures, or an increase in viscosity, but rather by the development of an elastic network across the condensates.

### 2.5 Relationship between driving force for phase separation and the optomechanical properties of condensates

To interrogate the link between the driving force for phase separation and optomechanical properties, we conducted QPI and Brillouin microscopy experiments with designed variants of the A1-LCD. The amino acid sequences can be found in the supplement (Table S 14). The variants differ in their composition of aromatic residues, which were previously identified as the main drivers of pairwise interactions that generate three-dimensional networks and mediate homotypic phase separation in this system [54]. By substituting the other aromatic residues with tryptophan, which has the largest aromatic ring system, we were able to increase the driving force for phase separation of the resulting protein variant. In the first mutant, only the 7 tyrosine residues of the wild-type (WT) were substituted with tryptophan residues (YtoW). In a second mutant, all 12 phenylalanine residues were substituted with tryptophan residues (FtoW), further increasing the phase separation propensity. Finally, all 12 phenylalanine and 7 tyrosine residues were mutated to tryptophan residues (allW) yielding a strongly phase-separating variant. Together, these four variants provided a broad range of phase separation propensities which is reflected in a range of saturation concentrations spanning two orders of magnitude [12]. We observed an overall negative trend of the saturation concentration with the protein concentration, but no consistent correlation (Figure 7A). Figure 7B shows that 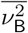 follows a logarithmic relationship with the saturation concentration according to 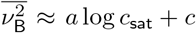. The Brillouin linewidth, on the other hand, shows similar values for allW and FtoW, as well as for YtoW and WT, but does not follow a consistent trend with the saturation concentration.

**Figure 7:**
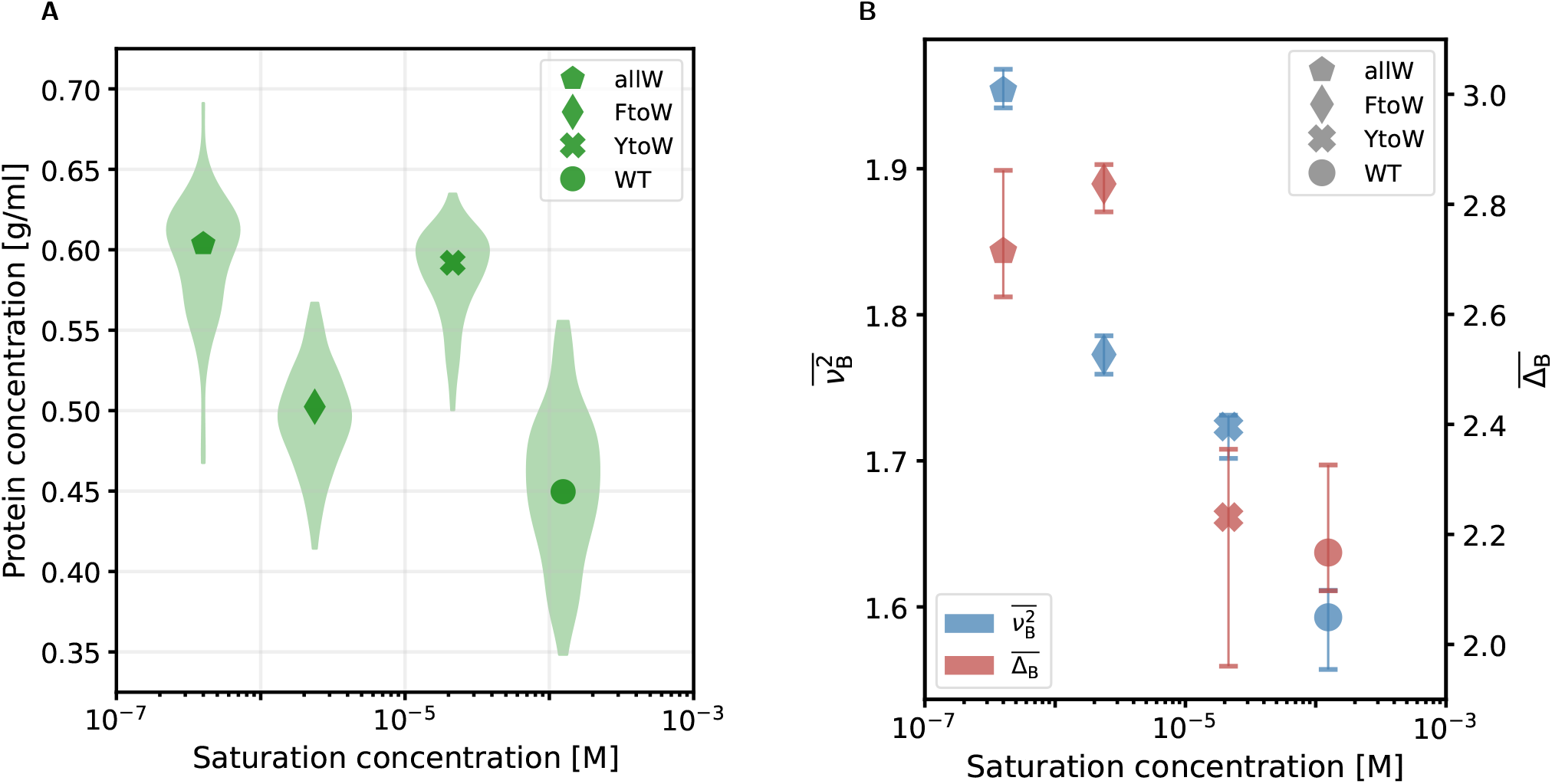
Physical properties of different A1-LCD-variants against their saturation concentration. Measurements were conducted at 23 °C and 150 mM NaCl. A: Protein concentration. Refractive index and volume fraction are shown in Figure S 10. B: Brillouin shift squared (left axis) and Brillouin linewidth (right axis) normalized to the value of the dilute phase (same value for all conditions). Markers indicate median values; error bars represent the range containing 68.3% of the data points that are closest to the median.

In summary, we find that altering pairwise protein-protein interaction strengths by changing the molecular grammar of protein sequences does not only change the saturation concentration but also the physical properties of the respective condensates. However, the presented data highlights that even though measured quantities are related to intermolecular interactions, they are not necessarily connected by simple relationships.

## 3 Discussion

In this work, we present Brillouin microscopy as a promising tool for the characterization of the physical properties of biomolecular condensates, as it introduces new ways to quantify the average molecular interaction and dissipation strength. The combination with QPI offers the possibility to obtain the condensate density and allows a precise calculation of the longitudinal modulus. In contrast to other commonly employed techniques, our experimental approach is non-invasive, does not rely on fluorescent labels, and can be applied *in vitro* and *in vivo*.

We showed that parameters with an impact on the phase diagram (i.e., temperature, ion concentration, and amino acid sequence) also affect the physical properties of protein condensates. Molecular interactions are therefore not only related to the saturation concentration but can be also accessed directly via the optomechanical properties of the dense phase. In general, the absolute values of longitudinal modulus and protein concentration are relatively high for condensed biological materials. We found protein concentrations between 0.26 g/ml for FUS condensates at 0.5 M KCl and 0.60 g/ml for A1-LCD allW and longitudinal moduli between 2.65 GPa for FUS condensates at 40 °C and 4.26 GPa for A1-LCD WT at 0.5 M NaCl. Typical dry mass densities for HeLa cells for example cover a range from about 0.07 g/ml in the nucleoplasm to 0.15 g/ml in nucleoli [55], corresponding to a longitudinal modulus of about 2.4 GPa to 2.5 GPa. Intracellular FUS condensates in living cells fall in the same range [31].

Similar values for protein concentration (0.26 g/ml) and longitudinal modulus (3.1 GPa) were previously found for intracellular polyQ aggregates in vivo [31]. Concentration measurements of phase-separated FUS-EGFP [56], as well as low-complexity domain molecules of FUS [57, 58] and A1-LCD [39, 54] found values of a few hundred mg/ml. Furthermore, a recent study employing QPI measured 0.34 g/ml for FUS-mEGFP [47] which is in line with our findings. Overall, these results show that the average molecular interaction strength and density of the investigated condensates *in vitro* are higher in living cells.

The differences between monocomponent condensates *in vitro* and complex multicomponent condensates in cells may underlie these substantial differences. Our study highlights the fact that the physical properties of protein condensates may be variable and context dependent, responding to external parameters such as thermodynamic variables. We also demonstrate how the physical properties can be tuned by altering the amino acid sequence. Therefore, the complex biochemical environment in living cells will modulate their properties, for example, due to the incorporation of additional molecules into the condensates or by post-translational modifications [59].

Uncertainties for the calculation of the density, protein concentration, and related quantities (i.e., protein volume fraction, longitudinal modulus) arise from the application of dextran as a crowding agent in some of the conditions studied (see Section S 8 for further details). It was found that in the absence of specific interactions and with high molecular weight crowders, depletion effects dominate and the crowder molecules are excluded from the condensates [60]. In light of these findings, the underlying assumption for our calculation of the density was that the crowder (dextran T500) is largely excluded from the protein condensates, and the values for *α* and *θ* were chosen accordingly. Nevertheless, we found that the addition of 5% dextran resulted in a small reduction in Brillouin shift and linewidth of the FUS condensates at low KCl concentrations (see Section S 9 for further details). Whether this difference arises from a partitioning of crowder molecules into the condensates still needs to be investigated. Furthermore, we assumed identical ion concentrations in the dilute and the dense phase, while we aware that ion partitioning in dilute and dense phase may differ [61]. The impact of this uneven ion distribution on the refractive computation, however, should be negligible due to the weak dependency of ion concentration on refractive index (see Section S 2 for further details).

It should be noted that refractive index measurements of non-spherical membraneless organelles in cells would require a tomographic imaging technique, such as ODT.

Our investigation of PEG solutions and protein condensates at different temperatures revealed characteristic behavior for different solute concentrations. We found that water dominates the response in the dilute phase of the phase separated protein samples and in PEG solutions up to a concentration of about 10%. For PEG concentrations higher than 20%, and in the investigated protein condensates, the polymers have a greater impact on viscoelasticity. This fits with the fact that both polymers have now exceeded their overlap concentrations and have entered the semi-dilute regime, greatly increasing the likelihood of interactions and entanglements [62]. Consequently, we find a decrease in molecular interaction and dissipation strength upon an increase in temperature. Such behavior is also known from glass-forming liquids [49, 63]. Consistent with the decrease in Brillouin linewidth, polymer solutions typically show lower viscosities at higher temperatures [46, 64]. A reduction in viscosity with increasing temperature was also recently observed in A1-LCD condensates [12].

The lowering of molecular interaction strength due to a temperature increase was also reflected in smaller protein concentrations in the condensates. Our findings are in line with experimental phase diagrams of A1-LCD [54] as well as coarse-grained simulations of FUS condensates [65]. We speculate that the local maximum of the Brillouin linewidth at 15 °C could be the hallmark that the molecular relaxation time in the A1-LCD condensates moved into the regime of the probed timescale of few hundred nanoseconds [26]. Such a transition was already observed for gelatin [49] at protein weight fractions above 0.4 (corresponding to volume fractions of about 0.3) and would be consistent with a further increase of protein volume fraction in A1-LCD condensates for temperatures below 22 °C. These results highlight that temperature strongly impacts the physical properties of protein condensates.

We also investigated the effect of varying ion concentrations on the physical properties of protein condensates, as ionic strength is a strong determinant of phase separation. In the case of FUS, the complex interplay of ion concentration and molecular interactions leads to reentrant phase separation [38]. At a low salt concentration below 125 mM, electrostatic interactions contribute to phase separation. These interactions are shielded by the presence of higher ion concentrations. At very high KCl concentrations above 1.5 M, however, hydrophobic and non-ionic interactions drive the system again into a phase-separating regime.

Our results agree with these interaction regimes and further reveal the ion dependence of molecular interactions in FUS condensates. In the presence of a crowding agent, we found a drop of protein concentration in the dense phase at KCl concentrations above 100 mM associated with a reduction of Brillouin shift and linewidth. Furthermore, we discovered that molecular interactions were strengthened at KCl concentrations above 1 M but did not reach the levels of the low salt conditions, even above the previously reported reentrant point of 1.5 M KCl. Also, protein concentration and volume fraction reached their maximum at the lowest ion concentrations. We conclude that the overall molecular interactions are weaker in the high salt regime. Apparently, the reduction of electrostatic interactions is not completely compensated by the hydrophobic and non-ionic interactions.

In the case of A1-LCD an increase of ionic strength leads to a reduction of the saturation concentration [50]. Note that fulllength hnRNPA1 shows the opposite behavior. As for the FUS condensates, we found that molecular interaction strength and density of the condensates correlated with the saturation concentration. Accordingly, a larger NaCl concentration led to higher protein concentration in the dense phase and a larger molecular interaction and dissipation strength.

Protein condensates are metastable fluids that can undergo changes in their morphologies and material properties. This may manifest in a slowing of molecular dynamics upon condensate aging or even in the formation of solid aggregates [66]. Although the transition of the majority of FUS condensates into fibrous aggregates was previously reported to take place after 12 h [6], we only observed Ostwald ripening even 36 h after phase separation. Our observations are in agreement with the cryo-EM images of Jawerth et al. [51], where morphological changes were only occasionally visible on the condensate interface but not in the bulk material. Along the same line, recent publications suggest that the aging of FUS protein condensates is an inhomogeneous process resulting in different material properties within the same condensate [67], and solidification can be promoted by the condensate interface [52].

We found that the aging process of the FUS condensates is accompanied by an increase of the Brillouin shift at low ion concentrations, while a change in linewidth and protein volume fraction was not detected. Our results may be interpreted as increasing polymer crosslinking, similar to what was found in gelatin [49] and gelatin methacryloyl (GelMA) [68]. The change of Brillouin shift and longitudinal modulus induced by crosslinking of a polymer solution is typically small if the polymer volume fraction is kept constant [49, 69, 70]. An increase in shear viscosity as reported previously [51] should have been reflected in a larger Brillouin linewidth, although the relationship between shear viscosity and Brillouin linewidth is not straightforward [71] (see also Section S 5).

Interestingly, the increase of the Brillouin shift was suppressed at a KCl concentration of 2.5 M, suggesting that a high abundance of ions attenuates the molecular interactions driving the aging process. A similar effect (i.e., suppression of aging) of high salt concentrations was previously reported for PGL-3 condensates [51]. These results suggest that the electrostatic interactions operating at low ion concentrations not only result in higher protein concentration and stronger average molecular interactions but are also important for the aging process of FUS condensates. The chemical crosslinking via the addition of glutaraldehyde led to a strong increase in the Brillouin shift and linewidth illustrating the effect of a complete condensate solidification.

We conclude that the ion-dependent aging that we observed was not the transition into semi-crystalline fibers, as this would have been associated with morphological changes, a large increase in Brillouin shift, and probably changes in Brillouin linewidth and protein volume fraction.

Our investigations of A1-LCD variants with increasingly strong driving forces for phase separation revealed an overall negative trend of saturation concentration with protein concentration and Brillouin linewidth of condensates, but not an absolute correlation. Specifically, YtoW condensates have a higher density than FtoW condensates despite having a higher saturation concentration and, the FtoW variant has a higher Brillouin linewidth than may be expected compared to the YtoW and allW variants. These two observations may be related to the length scales of the underlying interactions or the modulation of hydrodynamic effects by the different types of aromatic interactions. We have recently observed a similar decoupling of viscosities from the driving force for phase separation in variants that are dominated by different types of aromatic residues [12].

The negative trend with the saturation concentration is more apparent for the Brillouin shift; hence, the interaction strength between the proteins impacts both the saturation concentration and the average molecular interaction strength within the condensed phase. A comparison of A1-LCD and FUS shows that the FUS protein forms less dense condensates with weaker interactions compared to the A1-LCD variants, although the saturation concentration of FUS is relatively low (about 2 µM at 150 mM KCl) compared to most of the A1-LCD variants. One should note here that in the case of FUS, we studied the full-length protein with a GFP-tag, not the low-complexity domain in isolation. While we have not explicitly investigated the impact of a fluorescent label, the comparison of the properties of condensates assembled by labeled and unlabeled protein would be an interesting topic for future studies as many experimental techniques in the field of biomolecular condensates are based on the use of fluorescent labels. Our findings also further substantiate the sensitivity of the employed optical techniques to molecular interactions underlying phase separation. The sequence-dependent effects for protein concentration and Brillouin linewidth indicate an interesting decoupling of thermodynamics and hydrodynamics in variants that are dominated by different types of aromatic residues [12].

Finally, we want to discuss our methods and results with regard to biological systems. Over the past years, fluorescence microscopy has been the workhorse of molecular biology. It casts a spotlight on a selected molecular species, allows to localize them, and thereby enabled the discovery of membraneless organelles with liquid-like properties [72]. The majority of the abundant molecules in the investigated organelle and around remains invisible, though. While intracellular condensates appear as dense accumulations under a fluorescence microscope, it often remains unclear to what extent such accumulations of molecules really provide a different physical environment than the surroundings. The material properties of the condensates, however, have a crucial impact on the behavior of their content.

Less dense condensates with similar material properties as the surrounding intracellular space may interact as reaction sites [1]. Condensates exhibiting strong molecular interactions and high density can serve as a cellular mechanism to shut down biochemical reactions as a response to external stress or to enable cellular dormancy [3]. Moreover, recent research suggests that some condensates might also exert forces on other cellular organelles [9]. Therefore, cells need to regulate and sometimes actively change the physical properties of membraneless organelles, for example during germ cell development [4] or in DNA repair sites [11]. Our research also revealed that external stimuli, as well as intrinsic properties of the proteins, give rise to many subtle changes in the physical properties of the investigated condensates, which cannot be simply covered by qualitative terms such as “liquid-like” or “solid-like”.

Unraveling the implications of these diverse material properties and how cells control them will be a major task for the field of biomolecular condensates. This task requires the measurement of bulk physical properties in a quantitative, robust, and non-invasive way as reported here. The bulk quantities we measure, such as protein concentration, volume fraction, as well as sound velocity, and longitudinal modulus, may also be useful in refining coarse-grained simulations of condensates [39, 73, 74, 75] which can complement experimental results.

### 3.1 Outlook

While the presented experimental approach has great potential to advance our understanding of the physical properties of biomolecular condensates in general, we identified some opportunities for further improvement. A useful modification of the Brillouin microscope could be the employment of a spectrometer with a larger free spectral range. While the free spectral range is sufficient for most biological samples, we found that in some of the experimental conditions, an unambiguous evaluation of the Brillouin shift was obstructed by an overlap of the Brillouin peaks.

A powerful addition to the current instrumentation would be the possibility to detect shear waves as demonstrated by Kim et al. [76]. This would allow to directly measure the shear modulus, extract the bulk modulus from the longitudinal modulus, and derive an even more detailed physical description of the sample. Moreover, the detection of shear waves could be another modality to quantify a solidification process, because liquids are not permissive for shear wave propagation. Furthermore, faster data acquisition will be important for future studies targeting condensates in living cells. While the refractive index distribution of a full field of view covering multiple cells can be measured in about a second [55], the employed Brillouin microscope with its confocal point scanning approach (approximately 0.1 s to 1 s per pixel) poses a bottleneck as small and dynamic condensates might diffuse out of focus during the data acquisition. However, the acquisition speed of Brillouin microscopes is rapidly increasing and new technical innovations including full-field Brillouin microscopes are on their way [77].

Less dynamic and larger condensates, such as Balbiani and nuclear amyloid bodies, may therefore be promising next targets for Brillouin microscopy. There are a couple of burning questions regarding this type of condensates which urgently need fresh experimental concepts. What are the differences between physiological and pathological amyloid bodies? Why can cells dissolve the physiological ones? Could differences in the physical properties of physiological and pathological amyloid bodies explain this? We are convinced that the presented methods can be an important building block for future experiments that may answer these and other fundamental questions.

## 4 Materials and Methods

### 4.1 Protein purification

#### 4.1.1 FUS-GFP

FUS-GFP protein was expressed from insect cells using a baculovirus expression system [78]. Cells were lysed using an LM10 Microfluidizer (Microfluidics, USA) at 10 000 psi in lysis buffer containing 50 mM Tris-HCl pH 7.4, 1 M NaCl, 5% glycerol, 1 mM DTT, 33 U/ml of benzonase, and complete, EDTA-free Protease Inhibitor Cocktail (Merck). Lysed cells were centrifuged at 60 000 RCF for 1 h to separate soluble from insoluble fractions. The soluble fraction was bound to amylose resin (NEB, USA) for 1 h and washed with 14 column volumes of wash buffer containing 50 mM Tris-HCl pH 7.4, 1 M NaCl, 5% glycerol, 1 mM DTT. The protein was cleaved from the MBP by loading one column volume of the same buffer supplemented with 6 µg/ml 3C protease (produced in house), and incubating for 3 h. Protein was eluted in two column volumes of the same wash buffer, and the elution fraction was concentrated to 5 ml using an Amicon Ultra Centrifugal Filter Unit (30 kDa MWCO, Merck). The concentrated protein was applied to size exclusion chromatography using a HiLoad 16/600 Superdex 200 pg (Cytiva) column on the Akta Pure (Cytiva) FPLC system in 50 mM Tris-HCl pH 7.4, 750 mM KCl, 5% glycerol, 250 mM urea, and 1 mM DTT storage buffer. The protein was concentrated to 60 µM – 100 µM using an Amicon Ultra Centrifugal Filter Unit (30 kDa MWCO, Merck) and stored at −80 °C.

#### 4.1.2 hnRNPA1-LCD

A1-LCD construct design and purification were carried out as described previously [12, 39]. In brief, the A1-LCD sequence was cloned into pDEST17, with a TEV cleavage site to allow for His-tag removal. Proteins were expressed in BL21(DE3) cells overnight at 37 °C using autoinduction media. The protein was purified from the insoluble fraction using denaturing buffers and nickel-chelating resin. The tag was cleaved with His-tagged TEV protease, which was removed with a second nickel column. The proteins were further purified on a size exclusion column under denaturing conditions.

### 4.2 Brillouin microscopy

The Brillouin microscope was already described in detail elsewhere [79]. Briefly, a diode laser (TA pro 780, Toptica, Germany) was stabilized to the D2 transition of Rubidium 85 at 780.24 nm. In order to suppress amplified-spontaneous emission, the light was transmitted through a Fabry–Pérot interferometer in a two-pass configuration and guided on a Bragg-grating. The light was coupled to a Zeiss Axiovert 200 M microscope stand (Carl Zeiss, Germany) and directed to the sample through a 40x/0.95 air objective (Carl Zeiss, Germany). The backscattered light was collected in a confocal manner and analysed with a two-stage VIPA-spectrometer. We performed ca. 40-50 point measurements for several protein condensates for each condition. The number of evaluated condensates for each figure is given in the supplement (Table S 1). Dilute phase measurements consist of ca. 30 spectra from one location.

Data analysis was conducted with a custom software written in Python [80]. Brillouin spectra were fitted with two individual Lorentzian peaks in the case of dilute phase and the sum of four Lorentzian peaks in the case of protein condensates. The latter approach was necessary as in most cases also the Brillouin peaks of the dilute phase were contributing to the spectrum while all peaks were overlapping. In order to improve the quality of the fit, a measurement of the dilute phase was used to constrain the Lorentzian peaks correspondingly. Note that reported linewidths were broadened by the spectrometer and also by the distribution of scattering angles collected by the objective [81]. Temperature was controlled via a custom-built temperature stage [82].

### 4.3 Quantitative Phase Imaging (QPI)

QPI was performed with a previously described setup [31] which is also suitable for ODT. It is based on Mach-Zehnder interferometry, employing a frequency-doubled Nd-YAG laser with a wavelength of 532 nm (Torus, Laser Quantum, UK). A 40x/1.0 water-dipping objective lens (Carl Zeiss, Germany) was used for sample illumination. The light diffracted by the sample was collected with a 63x/1.2 water immersion objective (Carl Zeiss, Germany). For all temperatures above room temperature (23 °C), the sample temperature was set by two foil objective heaters (Thorlabs, USA), one for each objective, controlled by a TC200 temperature controller (Thorlabs, USA).

Phase images were computed from raw holograms that were acquired at a straight angle using qpimage [83]. From the phase image covering the full field of view, individual FUS condensates were segmented by a custom Python software [84]. The refractive index of the individual FUS-condensates was calculated from the segmented phase images by fitting a sphere using the Rytov approximation employing qpsphere [85, 86]. Only droplets with a diameter greater than 2 µm were considered for the evaluation.

### 4.4 Density and protein volume fraction computation

The mass density *ρ* was calculated with the following formula based on a binary mixture model,

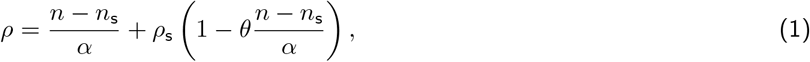

employing the assumption of volume additivity and the Biot mixing rule of refractive indices [79, 87]. In Equation 1, *n*_s_ denotes the refractive index of the solvent, *ρ*_s_ the density of the solvent, *θ* the proteins partial specific volume and *α* its refractive index increment. The first term in Equation 1 can be identified as the protein concentration *c* [88, 89]:

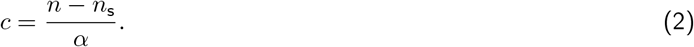

Using the information of the amino acid sequences of each protein, the values for *θ* and *α* were estimated based on the method proposed in [90] (see Table S 13). Here, instead of employing the Wiener mixing rule for dilute solutions under use in [90], the Biot mixing rule was used to maintain consistency with the above Equation 1 [87]. The refractive index of the solvent was measured with an Abbe-refractometer (2WAJ, Arcarda, Germany) (see Table S 11B and Figure S 1).

The protein volume fraction *ϕ* was calculated based on the protein concentration with the following formula:

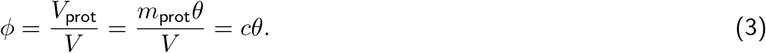

Effects of temperature on *α, θ*, and *ρ*_s_ were neglected and the respective values at room temperature were used to calculate refractive index, dry mass, volume fraction, and longitudinal modulus.

### 4.5 Longitudinal modulus computation

All measurements were performed in the back-scattering configuration, where the scattering angle *θ* (as defined in Figure 2) is approximately zero. Accordingly, the longitudinal modulus can be calculated by

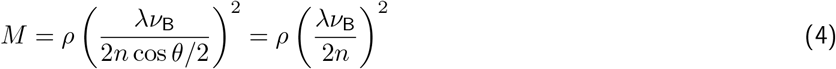

where *λ* denotes the incident wavelength of 780.24 nm. Equation 4 may be also written as

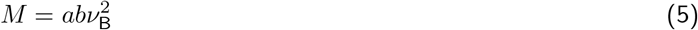

where 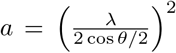 denotes a setup-specific constant and 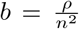 We found variations of *b* in the range of only about 2% (see Section S 6 for more details). Therefore, *b* ≈ const. and 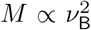 are reasonable approximations in the context of the presented data. For specimens with a more complex composition, where not only proteins are present, this approximation needs a more cautious treatment [87].

### 4.6 Data normalization

The values for the dilute phase, that were used to normalize longitudinal Modulus, Brillouin shift, and linewidth were either obtained by the respective median value of the dilute phase (Figure 3) or by taking the value corresponding value of a linear fit (Figure 4) in order to not transfer noise from the buffer measurement to the normalized values.

### 4.7 Fluorescence Recovery After Photobleaching (FRAP)

FRAP experiments were carried out with an Andor spinning disc microscope (Nikon TiE inverted stand, Nikon Apo 100x, NA 1.49 Oil objective, Andor iXon+ camera) equipped with a FRAPPA unit (Andor, Northern Ireland). Temperature was controlled the same way as for the Brillouin microscopy experiments. Condensates were bleached with a 488 nm pulse with 20 µs dwell time. Fluorescence recovery was recorded in a single focal plane. For each recorded temperature recovery curves of 5 individual condensates were recorded and analyzed with a custom python software using trackpy to correct for stage drifting [91].

### 4.8 Sample preparation

#### 4.8.1 FUS protein samples

If not stated differently, the protein stock solution (see Section 4.1.1 for further details) was diluted in a buffer containing 1 mM DTT, 20 mM PIPES at a pH of 7.4 to achieve a concentration of 5 µM FUS. If indicated, a concentration of 5%(w/w) dextran T500 (Pharmacosmos A/S, Denmark) was used to ensure phase separation at intermediate salt concentrations and high temperatures. For imaging the sample was loaded into a chamber, built up by a PEG-coated cover slide at the bottom and a silicone spacer (Grace Bio-Labs, USA) with a height of 0.9 mm and a diameter of 4.5 mm, resulting in a sample volume of about 14 µl. PEG coating was performed according to a protocol published earlier [92]. For Brillouin microscopy and FRAP experiments, the glass surface of the temperature control stage served as the upper boundary of the sample chamber. For the QPI measurements, an untreated cover slide was used instead, as the temperature control stage was not compatible with the QPI-setup.

Condensate fixation was performed by the addition of 0.05% glutaraldehyde after the initialization of phase separation by dilution of the stock protein solution with water.

#### 4.8.2 hnRNPA1-LCD protein samples

Amicon Ultra centrifugal filters (Merck, Germany) were filled with 440 µl of 1 M MES pH 5.5 buffer. 60 µl of approximately 500 µM protein in 4 M GdmHCl was added. Filters were centrifuged for 15 min at 21 000 g. Afterwards, the sample was filled up to the initial volume of 500 µl with MES buffer. The centrifugation step was repeated until a 1000-fold dilution of the initial GdmHCl buffer was achieved. After the last centrifugation step, the centrifugal filter was refilled with 20 mM HEPES pH 7.0, and the previous procedure was repeated until a 10,000-fold dilution of the MES buffer was achieved. Phase separation was induced by adding NaCl to a final concentration of 150 mM. Samples were loaded on a PEG-coated cover slide and enclosed by an imaging spacer (Grace Bio-Labs, USA).

#### 4.8.3 PEG 3350 solutions

Polyethylene Glycol (Sigma, USA) was dissolved in water to obtain a stock solution with a mass fraction of 50%. PEG solutions with a lower mass fraction were derived from the stock solution by dilution with water. Temperature control and sample mounting were conducted as for the FUS protein samples (see Section 4.8.1).

## Acknowledgements

We would like thank Felix Reichel for providing a stock solution of PEG and Anna Taubenberger for helpful discussions. Financial support from the Deutsche Forschungsgemeinschaft (SPP 2191-Molecular mechanisms of functional phase separation, grant agreement number 419138906 to SA and JG) and from the NIH (R01NS121114 to T.M.) is gratefully acknowledged. The coating of coverslips by PEG was supported by the European Unions Horizon 2020 research and innovation programs No. 953121 (project FLAMIN-GO) and TDSU Lab-on-a-chip systems at Max Planck Institute for the Science of Light

## Author contributions

The project was initiated by TB and JG; RS built the Brillouin microscope; KK built the QPI setup; PM developed essential parts of the QPI analysis software and helped with the implementation; LL purified the FUS protein; WB purified A1-LCD variants; RG coated the cover slides with PEG; TB conducted the experiments with help of LL, WB, KK, RS, TF, ML; CM computed refraction increment and partial specific volume of the proteins; TB analyzed and visualized the data; JG, TM and SA provided critical discussions and financial support; TB wrote the initial draft. TB edited the initial draft with help from all other authors.

## Supplement

### S 1 Number of evaluated samples

**Table S 1:** Number of evaluated condensates N in Figure 3B and 3D.

| Protein | FUS |  |  |  | hnRNPA1 |  |  |  |  |  |  |
| --- | --- | --- | --- | --- | --- | --- | --- | --- | --- | --- | --- |
| Temperature [°C] | 10 | 20 | 30 | 40 | 10 | 15 | 20 | 25 | 30 | 35 | 40 |
| N | 7 | 8 | 8 | 8 | 10 | 8 | 8 | 8 | 10 | 8 | 8 |

**Table S 2:** Number of evaluated condensates N in Figure 3C.

| Protein | FUS |  |  |  |  |  | hnRNPA1 |  |  |
| --- | --- | --- | --- | --- | --- | --- | --- | --- | --- |
| Temperature [°C] | 22.5 | 25 | 30 | 35 | 40 | 45 | 25 | 30 | 35 |
| N | 175 | 365 | 228 | 212 | 140 | 205 | 176 | 348 | 389 |

**Table S 3:** Number of evaluated condensates N in Figure 4A.

| Protein / salt | FUS / KCl |  |  |  |  |  |  | hnRNPA1 / NaCl |  |  |  |
| --- | --- | --- | --- | --- | --- | --- | --- | --- | --- | --- | --- |
| Salt conc. [M] | 0.066 | 0.1 | 0.15 | 0.5 | 1 | 1.7 | 2.13 | 0.1 | 0.15 | 0.3 | 0.5 |
| N | 262 | 112 | 175 | 62 | 76 | 52 | 130 | 148 | 176 | 324 | 453 |

**Table S 4:** Number of evaluated condensates N in Figure 4B.

| Protein / salt | FUS / KCl |  |  |  |  |  | hnRNPA1 / NaCl |  |  |  |
| --- | --- | --- | --- | --- | --- | --- | --- | --- | --- | --- |
| Salt conc. [M] | 0.066 | 0.15 | 0.5 | 1 | 1.7 | 2.13 | 0.1 | 0.15 | 0.3 | 0.5 |
| N | 10 | 31 | 12 | 30 | 31 | 29 | 15 | 10 | 15 | 12 |

**Table S 5:** Number of evaluated condensates N in Figure 5B.

| Time point [h] | 3 | 12 | 24 | 36 |
| --- | --- | --- | --- | --- |
| N | 524 | 556 | 489 | 433 |

**Table S 6:** Number of evaluated condensates N in Figure 6A and 6B.

| Condition | fixed | 0.15 M KCl 5 % dextran |  |  |  | 0.1 M KCl |  |  |  | 2.5 M KCl |  |  |  |
| --- | --- | --- | --- | --- | --- | --- | --- | --- | --- | --- | --- | --- | --- |
| Time point [h] | 0 | 0 | 12 | 24 | 36 | 0 | 12 | 24 | 36 | 0 | 12 | 24 | 36 |
| N | 11 | 31 | 29 | 36 | 35 | 30 | 31 | 28 | 29 | 32 | 30 | 31 | 32 |

**Table S 7:** Number of evaluated condensates N in Figure 7A.

| Variant | allW | FtoW | YtoW | WT |
| --- | --- | --- | --- | --- |
| N | 133 | 290 | 396 | 176 |

**Table S 8:**
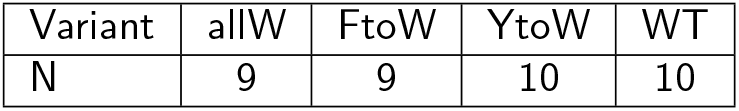
Number of evaluated condensates N in Figure 7B.

### S 2 Derivation of refractive index and density of buffer solutions

**Table S 9:** Refractive index and density of aqueous salt solutions. Refractive index values were measured with an Abbe-refractometer at room temperature (22 °C). Density values in (B) and (A) were taken from literature [1] and were measured at 25 °C. Density values in (C) were calculated via Equation 1, with *α*_Dextran_ = 0.14 ml/g [2] and *θ*_Dextran_ = 0.5959 ml/g [3]. A linear fit of the density values was used to approximate the density of the dilute phase for the calculation of the normalized longitudinal modulus in S 5B. The influence of PIPES / HEPES on the density was neglected.

| A |  |  | B |  |  | C |  |  |
| --- | --- | --- | --- | --- | --- | --- | --- | --- |
| H<sub>2</sub>O |  |  | H<sub>2</sub>O |  |  | H<sub>2</sub>O 5 % dextran |  |  |
| NaCl [M] | $n$ | $\rho$ [g/ml] | KCl [M] | $n$ | $\rho$ [g/ml] | KCl [M] | $n$ | $\rho$ [g/ml] |
| 0 | 1.334 | 0.997 | 0 | 1.334 | 0.997 | 0 | 1.341 | 1.0192 |
| 1 | 1.344 | 1.0361 | 0.5 | 1.339 | 1.01977 | 0.5 | 1.346 | 1.0393 |
| 2 | 1.353 | 1.0721 | 1 | 1.344 | 1.04134 | 1 | 1.351 | 1.0593 |
| 3 | 1.362 | 1.10557 | 1.5 | 1.3485 | 1.06189 | 1.7 | 1.3585 | 1.0874 |
| 4 | 1.37475 | 1.13684 | 2 | 1.3535 | 1.08152 | 2.13 | 1.362 | 1.1046 |
|  |  |  | 2.5 | 1.3585 | 1.10031 |  |  |  |

**Figure S 1:**
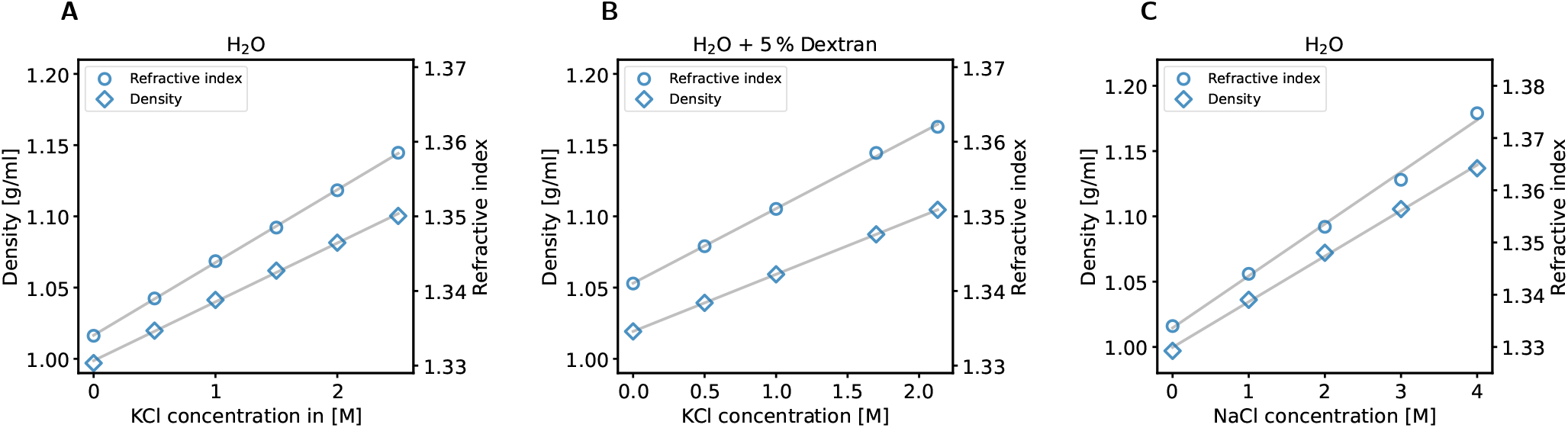
Density and refractive index of aqueous ion solutions. Values were taken from Table S 9. Refractive index and density for KCl dissolved in water (A), KCl dissolved in water with the addition 5% dextran (B), and NaCl dissolved in water. Fitting parameters for the linear fits are displayed in Table S 10.

**Table S 10:** Linear fitting parameters for refractive index *n* and density *ρ* of salt solutions. Parameters were calculated according to the equation *y* = *ax* + *b*, were *y* denotes either refractive index or density and *x* denotes the ion concentration.

| coefficient | $n_{\text{KCl}}$ | $\rho_{\text{KCl}}$ | $n_{\text{KCl+Dex}}$ | $\rho_{\text{KCl+Dex}}$ | $n_{\text{NaCl}}$ | $\rho_{\text{NaCl}}$ |
| --- | --- | --- | --- | --- | --- | --- |
| a | 0.0093/M | 0.0413 g/(ml M) | 0.0107/M | 0.0376 g/(ml M) | 0.0099/M | 0.0349 g/(ml M) |
| b | 1.3345 | 0.9987 g/ml | 1.3399 | 1.0218 g/ml | 1.3337 | 0.9997 g/ml |

**Table S 11:** Refractive index of buffer solutions. Refractive index values were measured with an Abbe-refractometer at room temperature (22 °C). A linear fit of the refractive index values was used to calculate protein concentration and volume fraction of the condensates.

| A<br>H<sub>2</sub>O + 20 mM HEPES |  | B<br>H<sub>2</sub>O + 20 mM PIPES |  | C<br>H<sub>2</sub>O + 20 mM PIPES + 5 % dextran |  |
| --- | --- | --- | --- | --- | --- |
| NaCl [M] | *n* | KCl [M] | *n* | KCl [M] | *n* |
| 0 | 1.3345 | 0 | 1.3345 | 0 | 1.3425 |
| 0.1 | 1.3355 | 0.15 | 1.3365 | 0.15 | 1.344 |
| 0.15 | 1.336 | 0.5 | 1.34 | 0.5 | 1.3465 |
| 0.3 | 1.3375 | 1 | 1.346 | 1 | 1.352 |
| 0.5 | 1.3395 | 1.7 | 1.35225 | 1.7 | 1.3595 |
| 1.5 | 1.349 | 2.13 | 1.356 | 2.13 | 1.3635 |
|  |  | 2.5 | 1.36 |  |  |

**Figure S 2:**
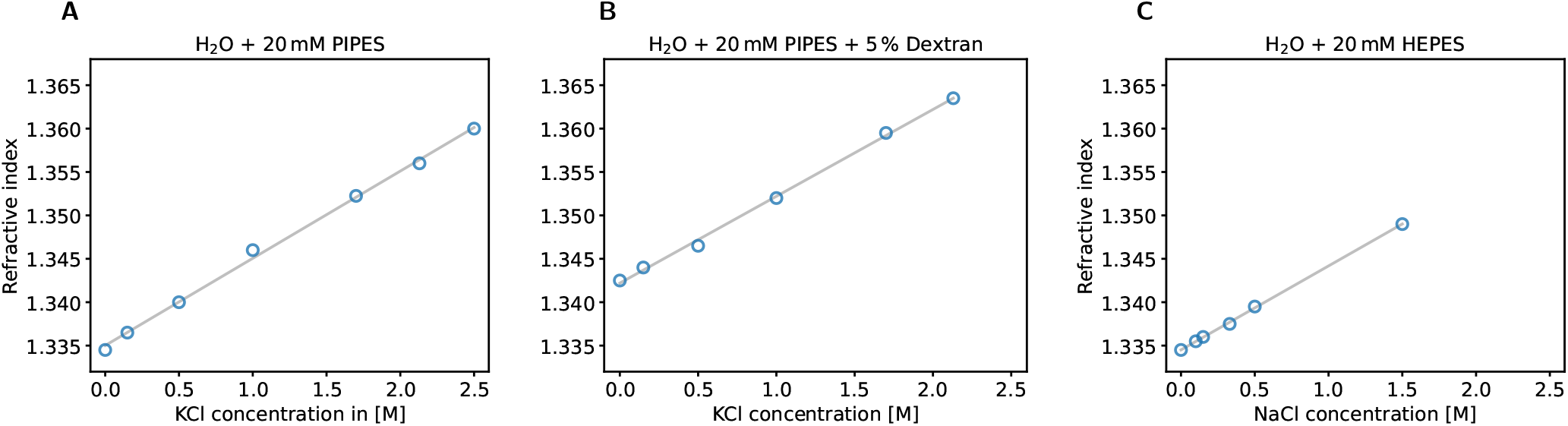
Refractive index of the buffer solutions. Values were taken from Table S 11. Refractive index for KCl dissolved in water (A), KCl dissolved in water with the addition 5% dextran (B), and NaCl dissolved in water(C). Fitting parameters for the linear fits are displayed in Table S 12.

**Table S 12:**
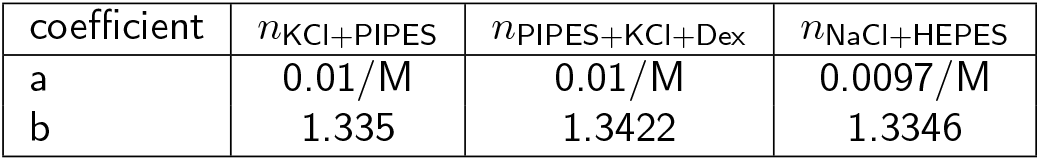
Linear fitting parameters for refractive index *n* of buffer solutions. Parameters were calculated according to the equation *y* = *ax* + *b*, were *y* denotes the refractive index.

### S 3 Refractive index increment and partial specific volume of the studied proteins

**Table S 13:**
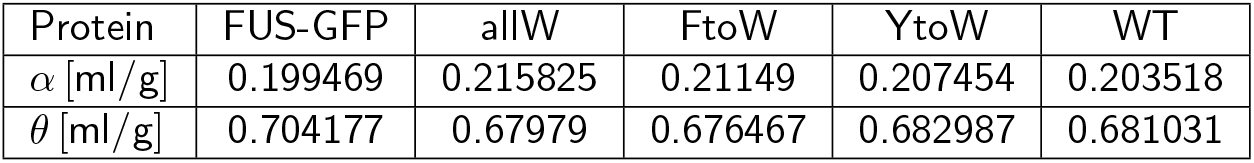
Refractive index increment *α* and partial specific volume *θ* for FUS-GFP and the investigated hnRNPA1 variants. Calculations were carried out as described in [4] based on the amino acid sequence.

### S 4 Brillouin spectra of hnRNPA1-LCD WT at different temperatures

Figure S 3 illustrates the challenges for the evaluation of the spectra of hnRNPA1-LCD WT recorded at low temperatures, with the employed Brillouin spectrometer. Because of the periodicity of the Brillouin spectrum, the correct calculation of the Brillouin shift relies on a clear attribution of each Brillouin peak to the elastic peaks (also called Rayleigh peaks) of the same scattering order (see also Figure 2, where a smaller fraction of a Brillouin spectrum is shown).

At a temperature of 20 °C, the Brillouin peaks corresponding to the protein condensates (the two peaks in the center around 7.59 GHz) are well separated. For larger Brillouin shifts or broader peaks (i.e., larger Brillouin linewidth), as for example in the spectra of Figure S 3C and Figure S 3B, the maxima of the peaks start to overlap.

**Figure S 3:**
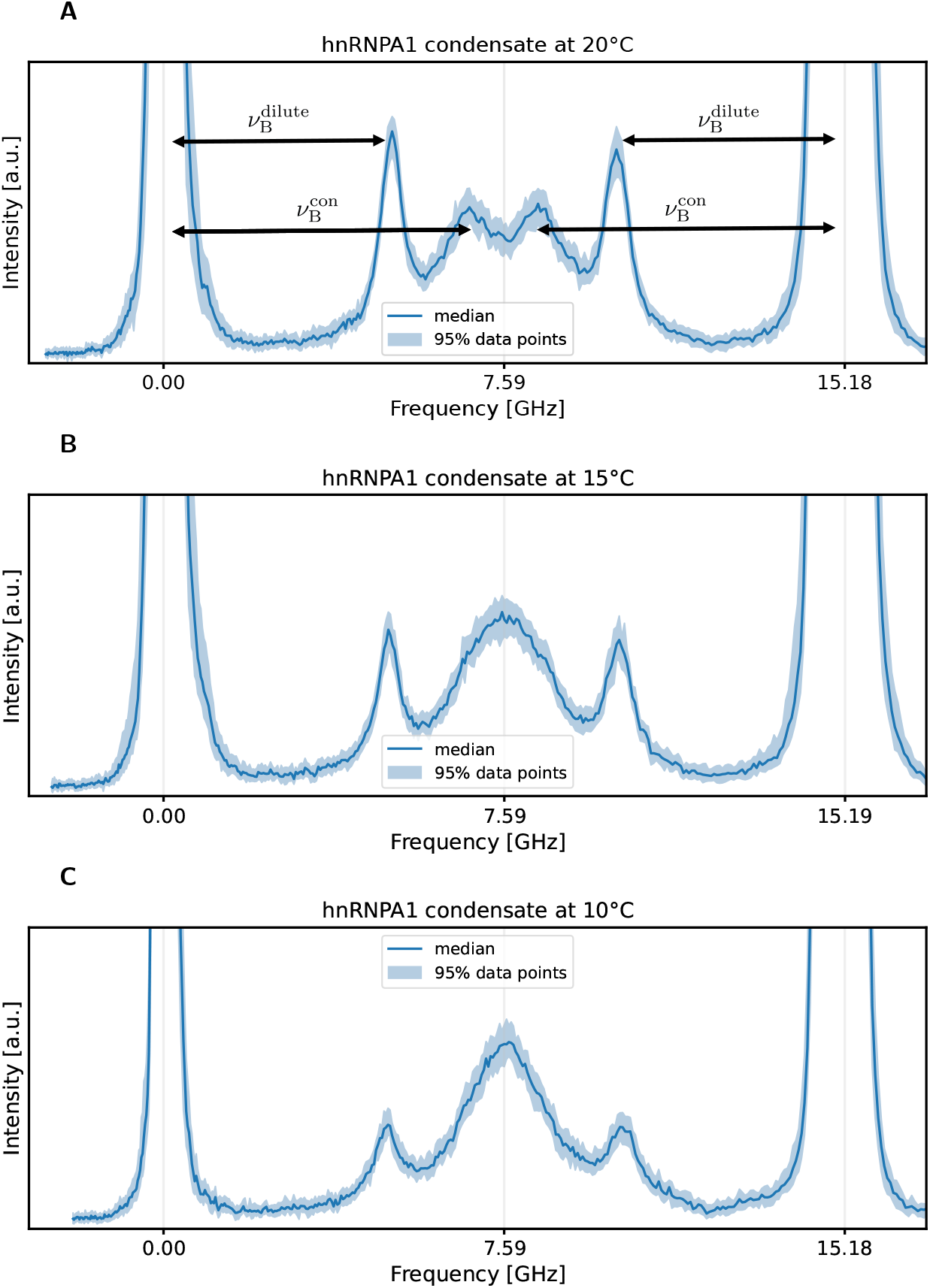
Brillouin spectra of hnRNPA1-LCD WT at different temperatures. Subfigures show spectra of three individual condensates at 20 °C (A), 15 °C (B) and 10 °C (C). Each dataset contains 50 individual spectra. The outer peaks (elastic peaks) correspond to different orders of the unshifted laser frequency, where their distance is given by the free spectral range of the spectrometer (approximately 15.2 GHz). The distance of a Brillouin peak to the corresponding elastic peak defines the Brillouin shift ν_B_. The recorded spectra contain four Brillouin peaks, where the outer two correspond to the dilute phase 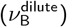, the inner ones to the dense phase 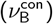.

While it is still possible to fit the overlapping peaks in such a condition, the attribution to the elastic peaks becomes ambiguous, in particular for the measurement at 10 °C. In other words, it is not clear weather the Brillouin peaks moved over the middle of the spectrum (corresponding to half of the free spectral range Δν_FSR_*/*2), resulting in a Brillouin shift greater than Δν_FSR_*/*2. The employed fitting procedure, however, assumes the Brillouin shift to be smaller than Δν_FSR_*/*2. Therefore, the calculated Brillouin shifts might be subjected to a systematic underestimation. As this potential underestimation does not affect the conclusions that were drawn from the results, we decided to keep the respective data in the figure and discuss the experimental challenges for these specific measurements. We want to point out here, that the Brillouin linewidth is not affected by this ambiguity. For comparison, Figure S 4 shows a modified version of Figure 3D assuming ν_B_ *>* Δν_FSR_*/*2 for hnRNPA1 at 10 °C. The issue of the overlapping peaks can be circumvented by the usage of a spectrometer with a greater free spectral range.

**Figure S 4:**
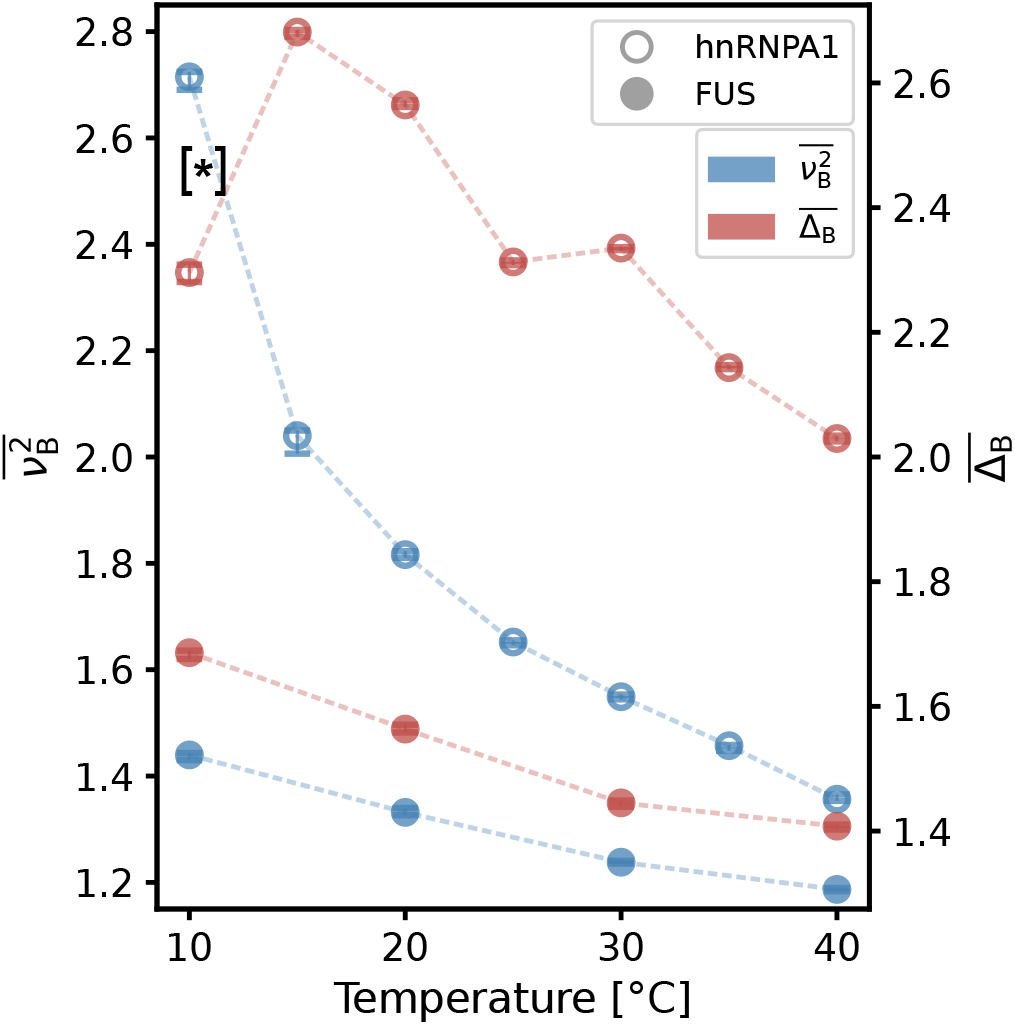
Modified version of Figure 3D. Reproduction of Figure 3D assuming ν_B_ *>* Δν_FSR_*/*2 for hnRNPA1 at 10 °C, indicated by [*], where Δν_FSR_ denotes the free spectral range. The modified value of the Brillouin shift 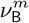 can be derived by the following formula: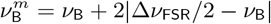.

### S 5 Relation of derived quantities with commonly used viscoelastic parameters

The most general description of the elastic properties of a material (in the linear regime) is the stiffness tensor in the generalized Hooke’s law with 21 independent components. All elastic moduli (bulk modulus *K*, longitudinal modulus *M*, Young’s modulus *E*, shear modulus *G*) are derived from the stiffness tensor and describe the material’s response for a specific type of load (e.g., shear stress). In the isotropic case, however, the stiffness tensor can be reduced to two independent parameters. Accordingly, any elastic modulus can be expressed by two other moduli. For the longitudinal modulus it is insightful to use bulk and shear modulus:

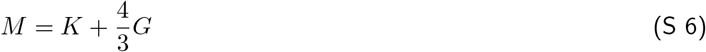

In the case of an ideal liquid *G* = 0 and *M* = *K*. While shear and Young’s modulus vanish (both describe a material deformation with constant volume), bulk and longitudinal modulus (both describe a material deformation with changing volume) are greater than zero. A deformation associated with a volume change requires a change in the equilibrium distance of the molecules (or atoms) assembling the material and are therefore directly coupled to molecular interaction strength. Protein condensates, in particular, are viscoelastic materials and can not be treated as simple liquids and consequently exhibit *G, E >* 0. Note that elastic properties, as well as viscous properties which are discussed in the subsequent text, may depend on the probed time scale (and accordingly frequency scale). A comprehensive overview of this topic is also provided in [5].

Assuming a Newtonian liquid, the Brillouin linewidth Δ_B_ in the employed backscattering geometry may be expressed by

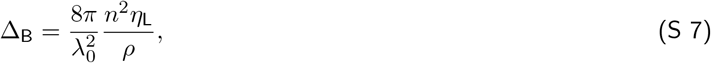

where *λ*_0_ is the laser wavelength (780.24 nm in our case) and *η*_L_ is the longitudinal viscosity [6]. The longitudinal viscosity in S 7 is defined analogue to the longitudinal modulus:

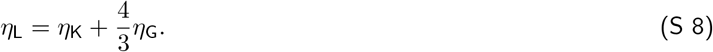

In fluid dynamics, *η*_L_ */ρ* is also known as kinematic viscosity. In the following section S 6 we find that 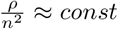. for the conducted experiments. In this case, it follows that

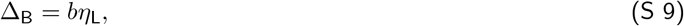

where *b* is an approximately constant value. It should be noted here that the linewidth is broadened by experimental setup, in particular due to the collection of a distribution of scattering angles which is defined by the objective’s numerical aperture [7]. Furthermore, there are additional wave decay mechanisms leading to an increase of Δ_B_, for example, due to static inhomogeneities [8], which are not included in this consideration. As the elastic moduli, the probed viscosity may depend on the rate at which it is measured (non-Newtonian behavior). Despite these caveats, it has been shown that the viscosity probed by a falling ball viscosimeter and viscosity calculated from Δ_B_ may exhibit a linear relationship for glycerol-water mixtures [6]. A detailed treatment of this topic can also be found in [9], where the relation of shear viscosity and Brillouin scattering derived viscosity of blood samples is investigated.

### S 6 Longitudinal modulus scaling

As already mentioned in the methods section 4.4 we used formula 1 to compute the mass density *ρ* = *ρ*(*n, α, θ*), which establishes a linear relationship with the refractive index *n*. Keeping the refraction increment *α* and the partial specific volume *θ* constant it follows the commonly used approximation 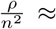. for small changes of *n*. Note that this approximation is only valid if the specimen exhibits constant values of *α* and *θ*. Proteins and lipids, for example, show different values for *α* and *θ*. Consequently, it has been shown that the assumption 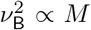 *M* is not valid for adipocytes [10], because proteins and lipids have very distinct values for *α* and *θ*. Figure S 5A illustrates the relative change of 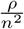 for the most extreme values of *n* appearing in this study using equation 1. We found changes of 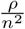 in the range of 2%. Due to this small change of 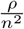 we found a linear correlation of 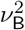 and *M* (Figure S 5B). Values for *n*_*s*_, *ρ*_*s*_ which were required for the calculation of the mass density based on Equation 1, were interpolated from the second and third column of Table S 11B, respectively.

**Figure S 5:**
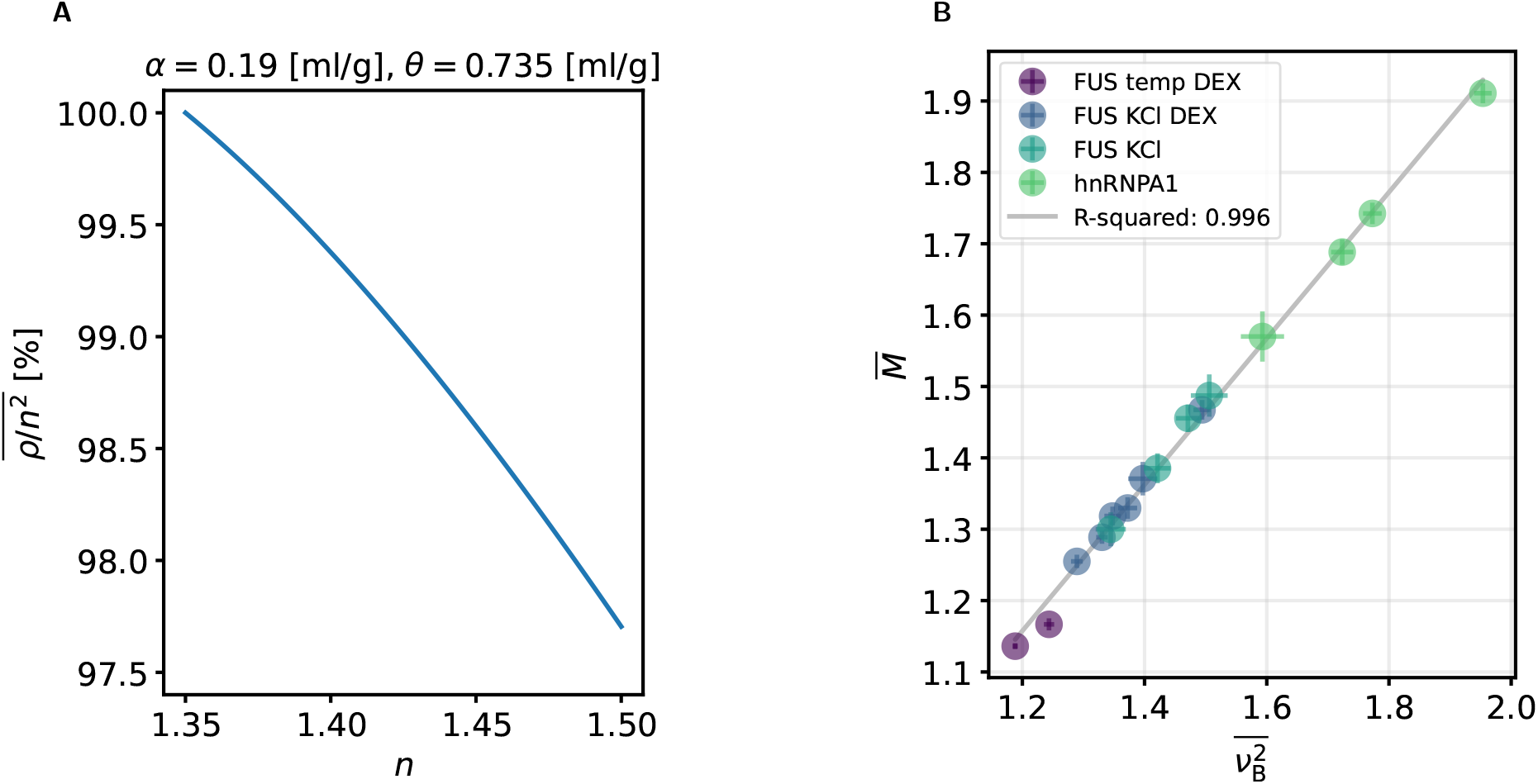
Impact of density and refractive index on longitudinal modulus computation. A: Relative changes of 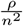 over the range of observed refractive index values. B: Correlation of normalized longitudinal Modulus with the normalized Brillouin shift squared. The presented values correspond to Figure 3D (FUS temp DEX), Figure 4B (FUS KCl DEX), Figure S 8B (FUS KCl) and Figure 7B (hnRNPA1), respectively. Error bars are based on Equation S 10 and the standard deviation of ν_B_.

The longitudinal modulus calculation was based on Brillouin shift, refractive index, refractive index increment, and partial specific volume, e.g., *M* = *M* (ν_B_, *n, α, θ*). The only contributions of random uncertainties originated from the measurements of Brillouin shift and refractive index. Based on the Gaussian error propagation we used the following formula to quantify the random uncertainties of the longitudinal modulus:

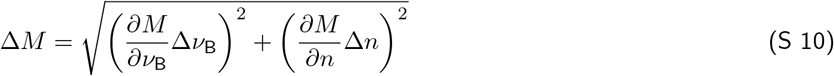

### S 7 Supplementary figures for Protein condensates at different ion concentrations

**Figure S 6:**
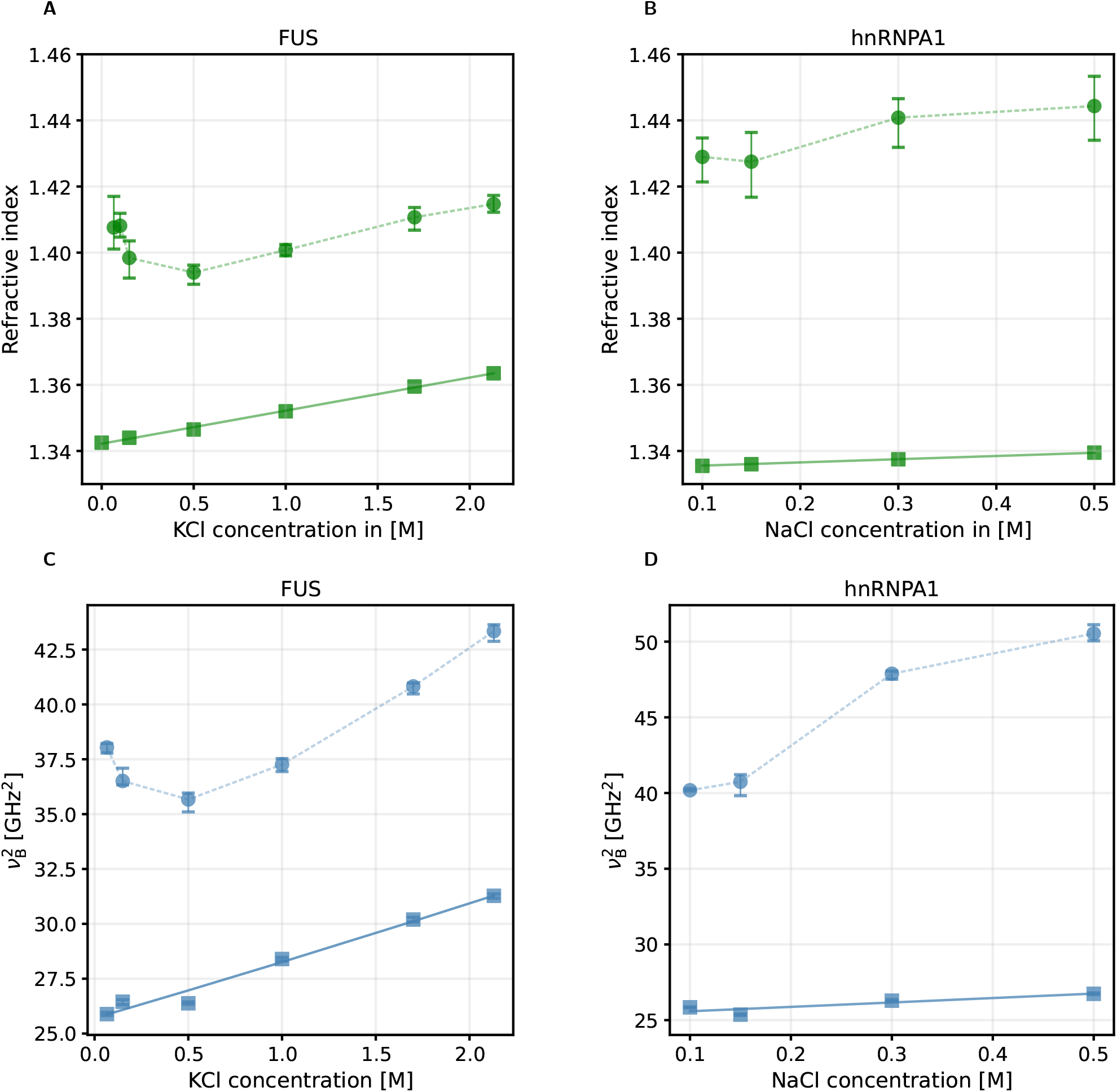
Comparison of buffer and protein condensates. Refractive index values of FUS condensates (circles) and buffer (squares) for various KCl concentrations (A) and for hnRNPA1 for various NaCl concentrations (B), respectively. Corresponding values of Brillouin shift squared of protein condensates (circles) and dilute phase (squares) for FUS (C) and hnRNPA1 (D), respectively. Markers indicate median values; error bars represent the range containing 68.3% of the data points that closest to the median.

### S 8 Uncertainties of the density computation

As already mentioned, deviations from the calculated density values may arise from the presence of dextran in the condensate (in case dextran was added to the buffer). The presence of dextran in the condensates would alter the effective values of refraction increment *α* and partial specific volume *θ* (i.e., *α*_Dextran_ ≈ 0.14 ml/g [2], *θ*_Dextran_ ≈ 0.6 ml/g [3]; *α*_protein_ ≈ 0.19 ml/g, *θ*_protein_ ≈ 0.735 ml/g [11]). The effect of these uncertainties on the calculation of the (dry) mass density is illustrated in Figure S 7.

Furthermore, ion concentration was assumed to be equal in dilute and dense phase. As the presence of ions impacts the refractive index, a non-uniform distribution of ions across the two phases would lead to an offset in the refractive index computation of the condensates. However, as this impact is relatively small (for KCl and NaCl in water we found d*n/* d*c*≤ 0.01 M^−1^, see Table S 12), we believe that this effect is likely negligible. Finally, the quantities *α* and *θ* might exhibit a certain variability upon changing ion and temperature conditions [12, 13, 14] that was not considered.

**Figure S 7:**
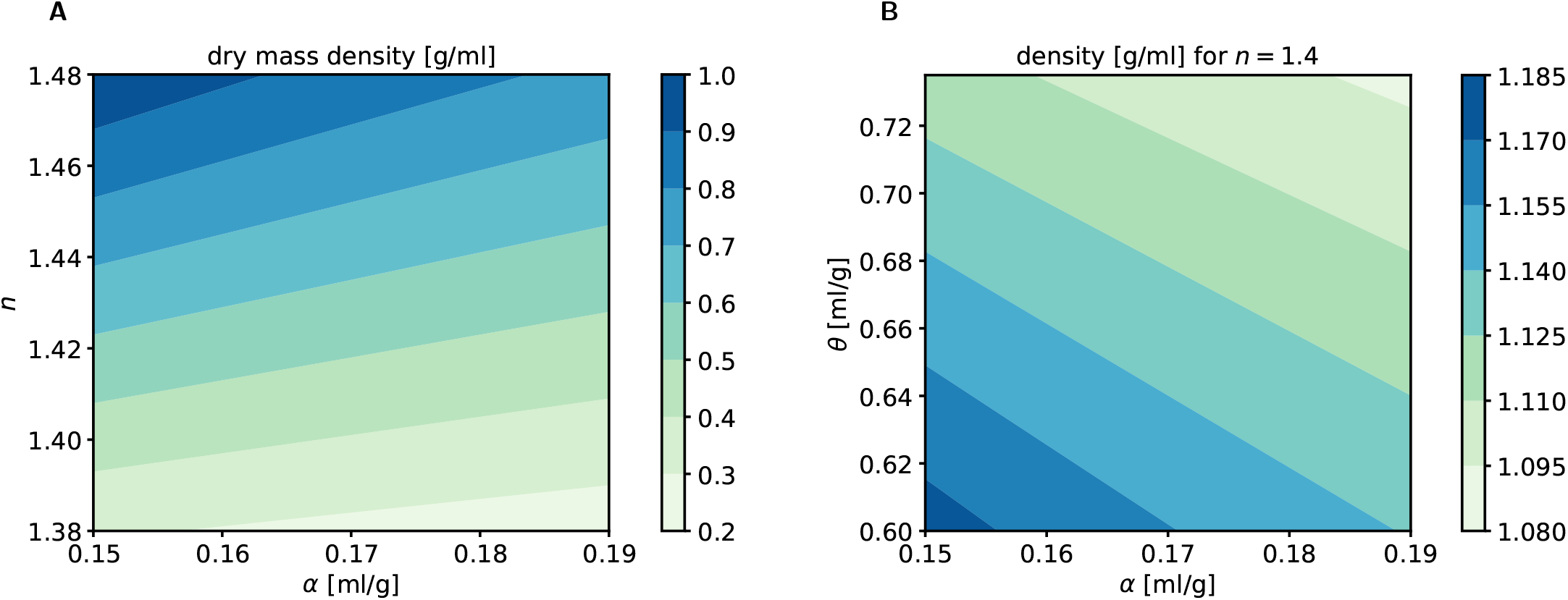
Impact of refractive index increment and partial specific volume on density computation. A: Solute concentration for different refractive index values and refraction increment. B: Mass density for mixtures of dextran and FUS for a given refractive index. Lower left of the plot corresponds to pure dextran, the upper right to pure FUS.

### S 9 Reference measurements of FUS condensates without dextran

In order to check the effect of dextran on the physical properties of FUS condensates we compared suitable conditions with and without the presence of dextran. While trends were qualitatively maintained, we found an offset due to the addition of dextran. In fact, we found that Brillouin shift and linewidth (Figure S 8), as well as refractive index (Figure S 9), protein concentration, and protein volume fraction of the FUS condensates were affected by the presence of a 5% dextran concentration. As phase separation was suppressed for temperatures above 30 °C and for intermediate ion concentrations, we could not acquire data in those regimes.

**Figure S 8:**
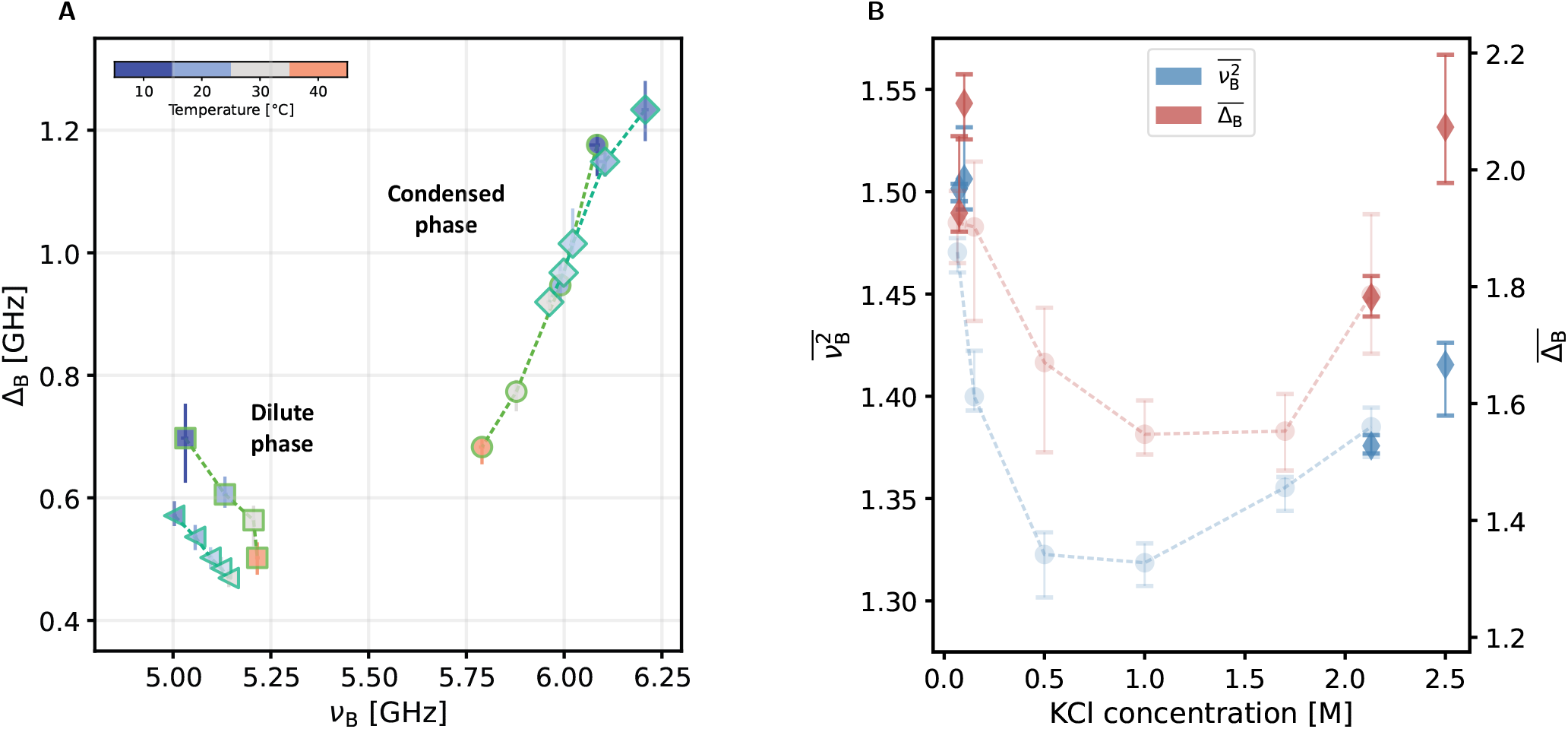
Evaluation of the impact of 5% dextran on Brillouin microscopy results. A: Values for Brillouin shift ν_B_ and linewidth Δ_B_ of dilute and condensed phase for various temperatures. The KCl concentration in was set to 150 mM.Triangles correspond to the dilute phase, diamonds show the condensed phase. Squares and circles correspond to the respective values with 5% dextran in the buffer as presented in the main text. The condensate markers contain values of 4 (15 °C), 5 (20 °C), 5 (25 °C), 5 (27.5 °C), and 5 (30 °C) individual condensates, respectively. B: Brillouin shift squared 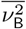 (left axis) and Brillouin linewidth 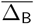 (right axis) of FUS condensates normalized to the respective values of the dilute phase (diamonds). Temperature was set to 20 °C. Circles correspond to the respective values with 5% dextran in the buffer as presented in the main text. The condensate markers contain values of 12 (75 mM), 30 (0.1 M), 12 (2.13 M), and 32 (2.5 M) individual condensates, respectively. Markers indicate median values; error bars represent the range containing 68.3% of the data points that are closest to the median.

**Figure S 9:**
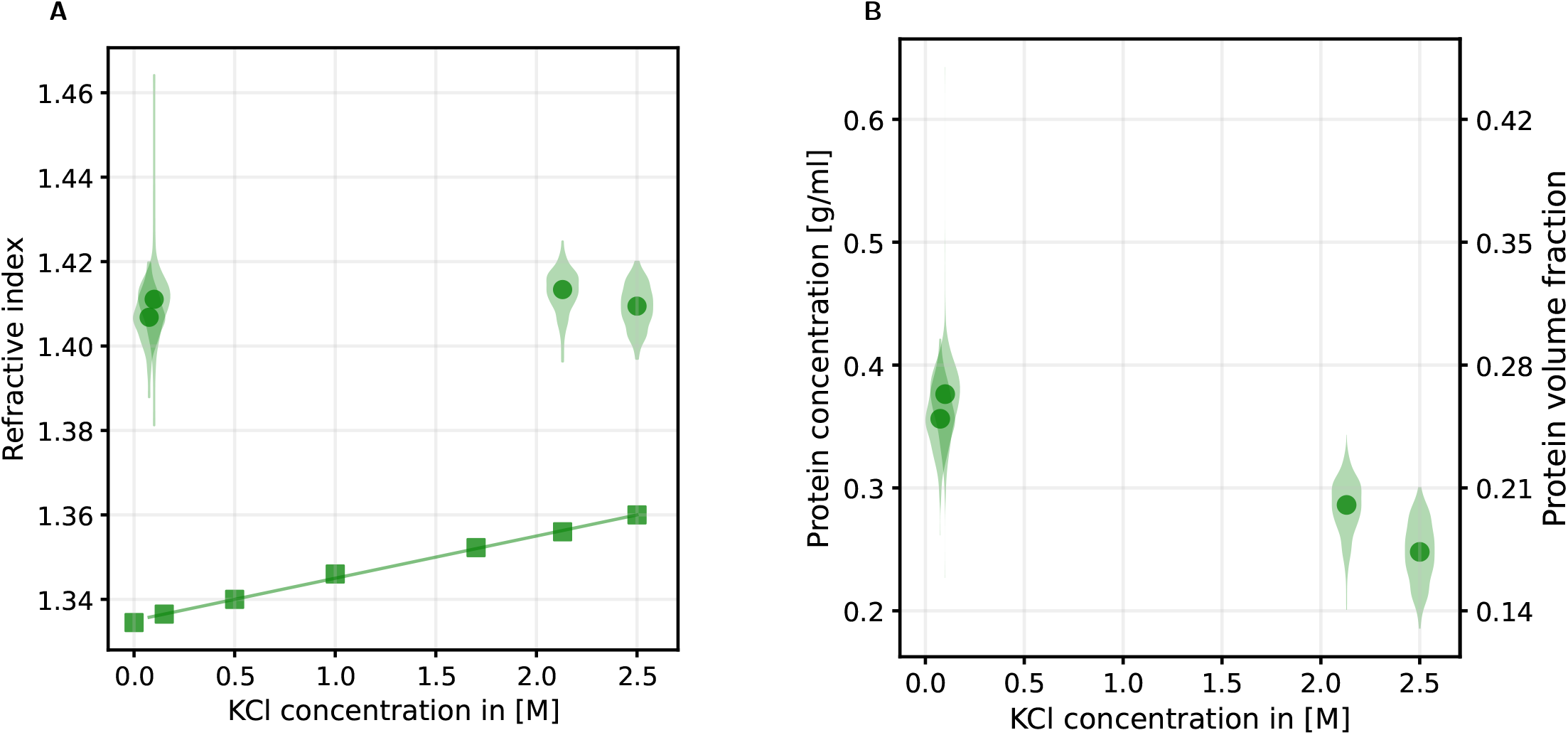
QPI measurements of FUS condensates. A: Refractive index of buffer and FUS protein condensates at different KCl concentrations. B: Corresponding protein concentration and protein volume fraction. Circles represent median values of FUS condensates, squares single measurements with a refractometer of the buffer solution. The data distributions contain values of 308 (75 mM), 408 (0.1 M), 561 (2.13 M), and 303 (2.5 M) individual condensates, respectively.

### S 10 Amino Acid sequences of hnRNPA1 variants

**Table S 14:**
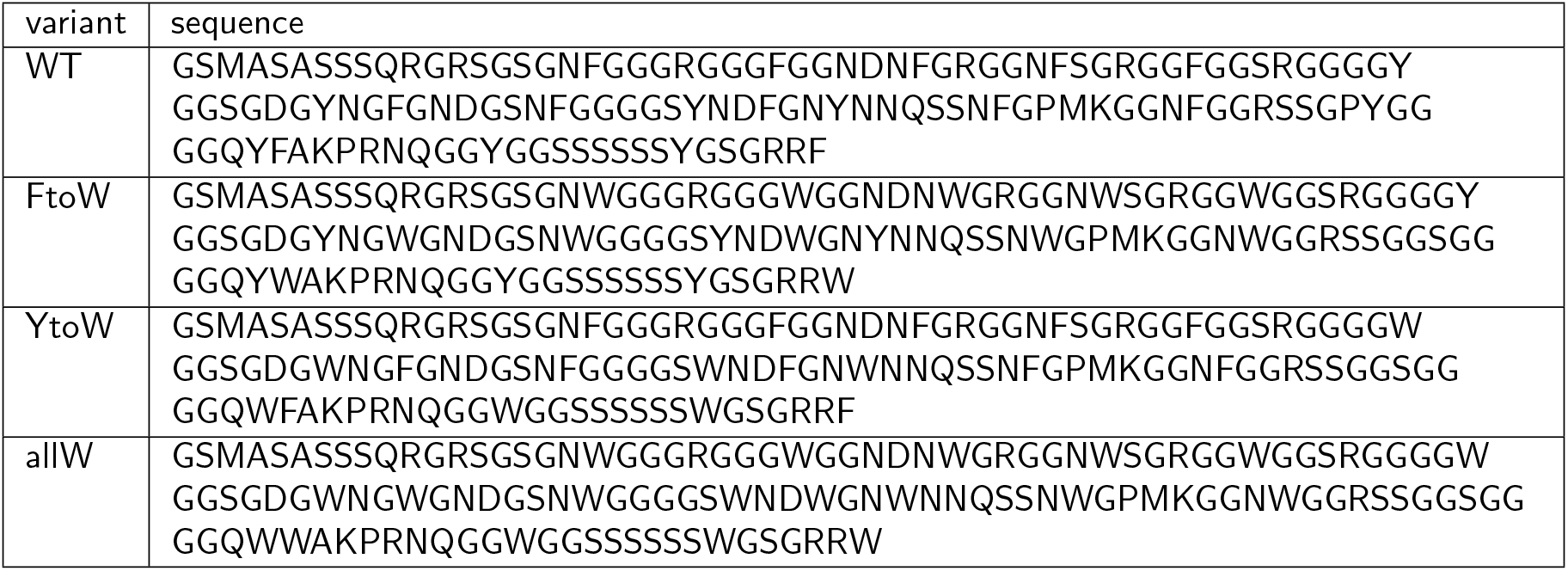
Amino acid sequences of the studied low complexity domains of hnRNPA1.

### S 11 Refractive index and protein volume fraction of hnRNPA1 condensates

**Figure S 10:**
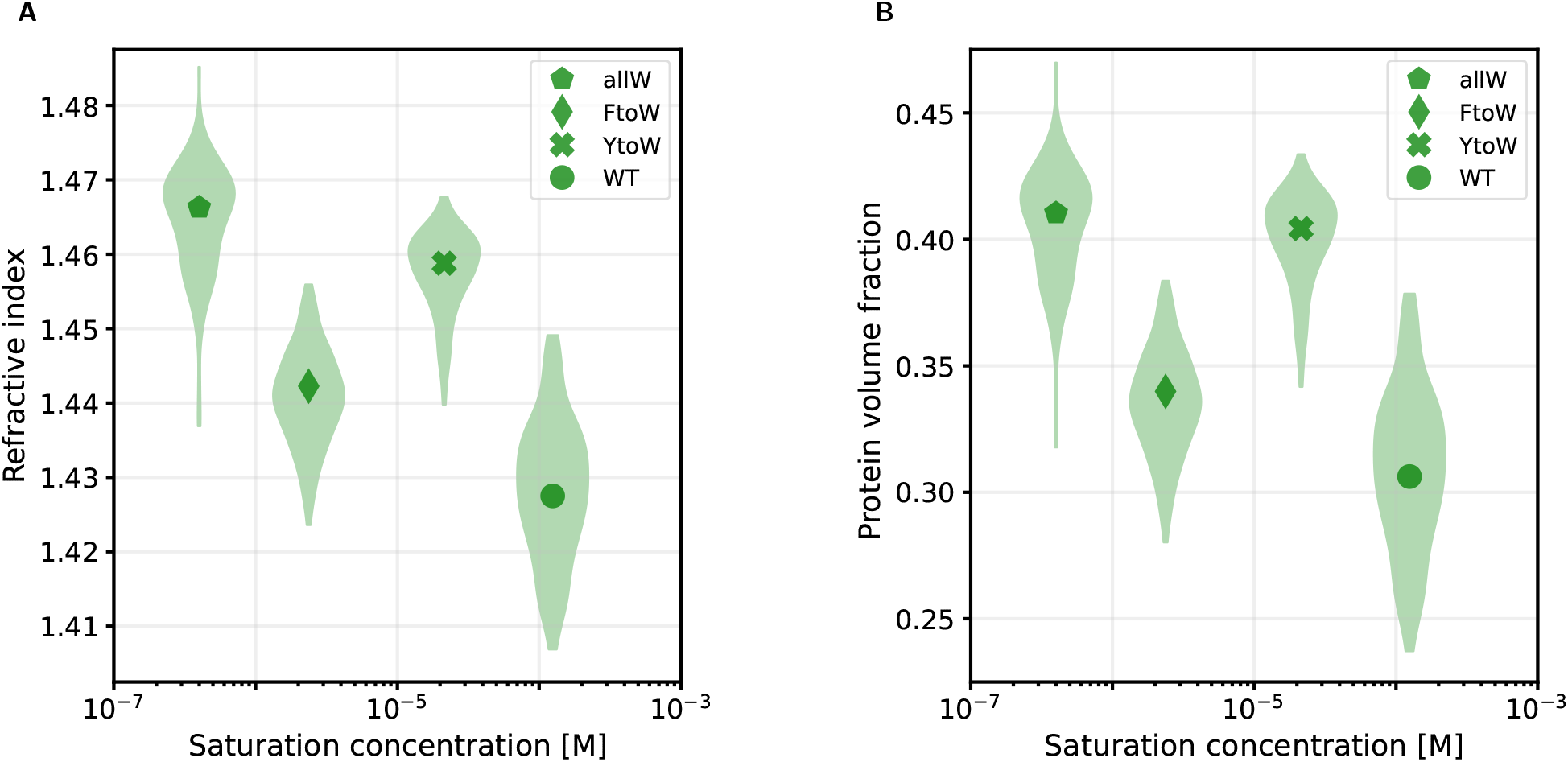
QPI measurements of hnRNPA1 condensates. Refractive index (A) and protein volume fraction (B) of hNRNPA1 condensates at 23 °C and 150 mM NaCl. Markers indicate median values.

## References

[1] Salman F. Banani et al. “Biomolecular condensates: Organizers of cellular biochemistry”. In: Nature Reviews Molecular Cell Biology 18.5 (2017), pp. 285–298. doi: 10.1038/nrm.2017.7.

[2] Clifford P. Brangwynne, Peter Tompa, and Rohit V. Pappu. “Polymer physics of intracellular phase transitions”. In: Nature Physics 11.11 (2015), pp. 899–904. doi: 10.1038/nphys3532.

[3] Jeffrey B. Woodruff, Anthony A. Hyman, and Elvan Boke. “Organization and Function of Non-dynamic Biomolecular Condensates”. In: Trends in Biochemical Sciences 43.2 (2018), pp. 81–94. doi: 10.1016/j.tibs.2017.11.005.

[4] Anne E. Dodson and Scott Kennedy. “Phase Separation in Germ Cells and Development”. In: Developmental Cell 55.1 (2020), pp. 4–17. doi: 10.1016/j.devcel.2020.09.004.

[5] Denis L.J. Lafontaine et al. “The nucleolus as a multiphase liquid condensate”. In: Nature Reviews Molecular Cell Biology 22.3 (2021), pp. 165–182. doi: 10.1038/s41580-020-0272-6.

[6] Avinash Patel et al. “A Liquid-to-Solid Phase Transition of the ALS Protein FUS Accelerated by Disease Mutation”. In: Cell 162.5 (2015), pp. 1066–1077. doi: 10.1016/j.cell.2015.07.047.

[7] Matthias Altmeyer et al. “Liquid demixing of intrinsically disordered proteins is seeded by poly(ADP-ribose)”. In: Nature Communications 6.1 (2015), p. 8088. doi: 10.1038/ncomms9088.

[8] Elvan Boke et al. “Amyloid-like Self-Assembly of a Cellular Compartment”. In: Cell 166.3 (2016), pp. 637–650. doi: 10.1016/j.cell.2016.06.051.

[9] Felipe Garcia Quiroz et al. “Liquid-liquid phase separation drives skin barrier formation”. In: Science 367.6483 (2020). doi: 10.1126/science.aax9554.

[10] Shana Elbaum-Garfinkle et al. “The disordered P granule protein LAF-1 drives phase separation into droplets with tunable viscosity and dynamics”. In: Proceedings of the National Academy of Sciences 112.23 (2015), pp. 7189–7194. doi: 10.1073/pnas.1504822112.

[11] Nagaraja Chappidi et al. “PARP1-DNA co-condensation drives DNA repair site assembly to prevent disjunction of broken DNA ends”. In: Cell 187.4 (2024), 945–961.e18. doi: 10.1016/j.cell.2024.01.015.

[12] Ibraheem Alshareedah et al. “Sequence-specific interactions determine viscoelasticity and ageing dynamics of protein condensates”. In: Nature Physics 20.9 (2024), pp. 1482–1491. doi: 10.1038/s41567-024-02558-1.

[13] Amandine Molliex et al. “Phase Separation by Low Complexity Domains Promotes Stress Granule Assembly and Drives Pathological Fibrillization”. In: Cell 163.1 (2015), pp. 123–133. doi: 10.1016/j.cell.2015.09.015.

[14] Yun R. Li et al. “Stress granules as crucibles of ALS pathogenesis”. In: Journal of Cell Biology 201.3 (2013), pp. 361–372. doi: 10.1083/jcb.201302044.

[15] Hong Joo Kim et al. “Mutations in prion-like domains in hnRNPA2B1 and hnRNPA1 cause multisystem proteinopathy and ALS”. In: Nature 495.7442 (2013), pp. 467–473. doi: 10.1038/nature11922.

[16] Anastasia C. Murthy and Nicolas L. Fawzi. “The (un)structural biology of biomolecular liquid-liquid phase separation using NMR spectroscopy”. In: Journal of Biological Chemistry 295.8 (2020), pp. 2375–2384. doi: 10.1074/jbc.REV119.009847.

[17] W Russ Algar et al. “FRET as a biomolecular research tool — understanding its potential while avoiding pitfalls”. In: Nature Methods 16.9 (2019), pp. 815–829. doi: 10.1038/s41592-019-0530-8.

[18] Ibraheem Alshareedah, Taranpreet Kaur, and Priya R. Banerjee. “Chapter Six - Methods for characterizing the material properties of biomolecular condensates”. In: Liquid-Liquid Phase Coexistence and Membraneless Organelles. Ed. by Christine D. Keating. Vol. 646. Methods in Enzymology. Academic Press, 2021, pp. 143–183. doi: 10.1016/bs.mie.2020.06.009.

[19] Nicole O. Taylor et al. “Quantifying Dynamics in Phase-Separated Condensates Using Fluorescence Recovery after Photobleaching”. In: Biophysical Journal 117 (7 2019). doi: 10.1016/j.bpj.2019.08.030.

[20] Alfredo Vidal Ceballos, Charles J. McDonald, and Shana Elbaum-Garfinkle. “Chapter Two - Methods and Strategies to Quantify Phase Separation of Disordered Proteins”. In: Intrinsically Disordered Proteins. Ed. by Elizabeth Rhoades. Vol. 611. Methods in Enzymology. Academic Press, 2018, pp. 31–50. doi: 10.1016/bs.mie.2018.09.037.

[21] Ibraheem Alshareedah et al. “Programmable viscoelasticity in protein-RNA condensates with disordered sticker-spacer polypeptides”. In: Nature Communications 12.1 (2021), p. 6620. doi: 10.1038/s41467-021-26733-7.

[22] Louise M. Jawerth et al. “Salt-Dependent Rheology and Surface Tension of Protein Condensates Using Optical Traps”. In: 121 (25 2018), p. 258101. doi: 10.1103/PhysRevLett.121.258101.

[23] Rachel S. Fisher and Shana Elbaum-Garfinkle. “Tunable multiphase dynamics of arginine and lysine liquid condensates”. In: Nature Communications 11.1 (2020), p. 4628. doi: 10.1038/s41467-020-18224-y.

[24] Brillouin, Léon. “Diffusion de la lumière et des rayons X par un corps transparent homogène - Influence de l’agitation thermique”. In: Ann. Phys. 9.17 (1922), pp. 88–122. doi: 10.1051/anphys/192209170088.

[25] LI Mandelstam. “Light scattering by inhomogeneous media”. In: Zh. Russ. Fiz-Khim. Ova 58.381 (1926), p. 146.

[26] Robert Prevedel et al. “Brillouin microscopy: an emerging tool for mechanobiology”. In: Nature Methods 16.10 (2019), pp. 969–977. doi: 10.1038/s41592-019-0543-3.

[27] Maurizio Mattarelli, Massimo Vassalli, and Silvia Caponi. “Relevant Length Scales in Brillouin Imaging of Bio-materials: The Interplay between Phonons Propagation and Light Focalization”. In: ACS Photonics (2020). doi: 10.1021/acsphotonics.0c00801.

[28] Irina Kabakova et al. “Brillouin microscopy”. In: Nature Reviews Methods Primers 4.1 (2024), p. 8. doi: 10.1038/s43586-023-00286-z.

[29] Y.P. Singh and R.P. Singh. “Compatibility studies on solutions of polymer blends by viscometric and ultrasonic techniques”. In: European Polymer Journal 19.6 (1983), pp. 535–541. doi: 10.1016/0014-3057(83)90206-9.

[30] Vidhya Jadhav et al. “Volumetric and compressibility studies and phase equilibria of aqueous biphasic systems of alcohols using phase diagram”. In: SN Applied Sciences 1.7 (2019), pp. 1–12. doi: 10.1007/s42452-019-0688-9.

[31] Raimund Schüßler et al. “Correlative all-optical quantification of mass density and mechanics of subcellular compartments with fluorescence specificity”. In: eLife 11 (2022). Ed. by Rohit V Pappu and Anna Akhmanova, e68490. doi: 10.7554/eLife.68490.

[32] Kyoohyun Kim et al. “Real-time visualization of 3-D dynamic microscopic objects using optical diffraction tomography”. In: Opt. Express 21.26 (2013), pp. 32269–32278. doi: 10.1364/OE.21.032269.

[33] Mirjam Schürmann et al. “Cell nuclei have lower refractive index and mass density than cytoplasm”. In: Journal of Biophotonics 9.10 (2016), pp. 1068–1076. doi: 10.1002/jbio.201500273.

[34] Giuseppe Antonacci and Sietse Braakman. “Biomechanics of subcellular structures by non-invasive Brillouin microscopy”. In: Scientific Reports 6.37217 (2016). doi: 10.1038/srep37217.

[35] Giuseppe Antonacci et al. “Background-deflection Brillouin microscopy reveals altered biomechanics of intracellular stress granules by ALS protein FUS”. In: Communications Biology 1.1 (2018), p. 139. doi: 10.1038/s42003-018-0148-x.

[36] Takuya Yoshizawa et al. “Nuclear Import Receptor Inhibits Phase Separation of FUS through Binding to Multiple Sites”. In: Cell 173.3 (2018), 693–705.e22. doi: 10.1016/j.cell.2018.03.003.

[37] Shujie Li et al. “Pressure and Temperature Phase Diagram for Liquid–Liquid Phase Separation of the RNA-Binding Protein Fused in Sarcoma”. In: The Journal of Physical Chemistry B 125.25 (2021), pp. 6821–6829. doi: 10.1021/acs.jpcb.1c01451.

[38] Georg Krainer et al. “Reentrant liquid condensate phase of proteins is stabilized by hydrophobic and non-ionic interactions”. In: Nature Communications 12.1 (2021), p. 1085. doi: 10.1038/s41467-021-21181-9.

[39] Anne Bremer et al. “Deciphering how naturally occurring sequence features impact the phase behaviours of disordered prion-like domains”. In: Nature Chemistry 14.2 (2022), pp. 196–207. doi: 10.1038/s41557-021-00840-w.

[40] M. Pochylski et al. “Structuring effects and hydration phenomena in poly(ethylene glycol)/water mixtures investigated by brillouin scattering”. In: Journal of Physical Chemistry B 110.41 (2006), pp. 20533–20539. doi: 10.1021/jp0620973.

[41] T.J.C. Hosea, S.C. Ng, and C.G. Oates. “A Brillouin scattering study of the glass transition in sucrose”. In: Food Hydrocolloids 4.2 (1990), pp. 137–147. doi: 10.1016/S0268-005X(09)80014-3.

[42] Frank J. Millero, Richard W. Curry, and Walter Drost-Hansen. “Isothermal compressibility of water at various temperatures”. In: Journal of Chemical & Engineering Data 14.4 (1969), pp. 422–425. doi: 10.1021/je60043a018.

[43] S. C. Santucci et al. “Is there any fast sound in water?” In: Physical Review Letters 97.22 (2006), pp. 1–4. doi: 10.1103/PhysRevLett.97.225701.

[44] Zdeňka Kolská et al. “Refractometric study of systems water-poly(ethylene glycol) for preparation and characterization of Au nanoparticles dispersion”. In: Arabian Journal of Chemistry 12.8 (2019), pp. 5019–5027. doi: 10.1016/j.arabjc.2016.11.006.

[45] Pedro Gonzalez-Tello, Fernando Camacho, and Gabriel Blazquez. “Density and Viscosity of Concentrated Aqueous Solutions of Polyethylene Glycol”. In: Journal of Chemical & Engineering Data 39.3 (1994), pp. 611–614. doi: 10.1021/je00015a050.

[46] Sharmin Ebrahimi and Rahmat Sadeghi. “Density, Speed of Sound, and Viscosity of Some Binary and Ternary Aqueous Polymer Solutions at Different Temperatures”. In: Journal of Chemical Engineering Data 60.11 (2015), pp. 3132–3147. doi: 10.1021/acs.jced.5b00290.

[47] Patrick M McCall et al. “Label-free composition determination for biomolecular condensates with an arbitrarily large number of components”. In: bioRxiv (2023), p. 2020. 10.25.352823.

[48] Giuseppe Antonacci et al. “Recent progress and current opinions in Brillouin microscopy for life science applications”. In: Biophysical Reviews 12.3 (2020), pp. 615–624. doi: 10.1007/s12551-020-00701-9.

[49] Michelle Bailey et al. “Viscoelastic properties of biopolymer hydrogels determined by Brillouin spectroscopy: A probe of tissue micromechanics”. In: Science Advances 6.44 (2020), eabc1937. doi: 10.1126/sciadv.abc1937.

[50] Erik W. Martin et al. “Interplay of folded domains and the disordered low-complexity domain in mediating hnRNPA1 phase separation”. In: Nucleic Acids Research 49.5 (2021), pp. 2931–2945. doi: 10.1093/nar/gkab063.

[51] Louise Jawerth et al. “Protein condensates as aging Maxwell fluids”. In: Science 370.6522 (2020), pp. 1317–1323. doi: 10.1126/science.aaw4951.

[52] Yi Shen et al. “The liquid-to-solid transition of FUS is promoted by the condensate surface”. In: Proceedings of the National Academy 120.33 (2023), e2301366120. doi: 10.1073/pnas.2301366120.

[53] Miloš Nikolić and Giuliano Scarcelli. “Long-term Brillouin imaging of live cells with reduced absorption-mediated damage at 660nm wavelength”. In: Biomedical Optics Express 10.4 (2019), p. 1567. doi: 10.1364/BOE.10.001567.

[54] Erik W Martin et al. “Valence and patterning of aromatic residues determine the phase behavior of prion-like domains”. In: Science 367.6478 (2020), pp. 694–699. doi: 10.1126/science.aaw8653.

[55] Kyoohyun Kim and Jochen Guck. “The Relative Densities of Cytoplasm and Nuclear Compartments Are Robust against Strong Perturbation”. In: Biophysical Journal 119.10 (2020), pp. 1946–1957. doi: 10.1016/j.bpj.2020.08.044.

[56] Jonas Ahlers et al. “The key role of solvent in condensation: Mapping water in liquid-liquid phase-separated FUS”. In: Biophysical Journal 120.7 (2021), pp. 1266–1275. doi: 10.1016/j.bpj.2021.01.019.

[57] Kathleen A. Burke et al. “Residue-by-Residue View of In Vitro FUS Granules that Bind the C-Terminal Domain of RNA Polymerase II”. In: Molecular Cell 60.2 (2015), pp. 231–241. doi: 10.1016/j.molcel.2015.09.006.

[58] Anastasia C. Murthy et al. “Molecular interactions underlying liquid-liquid phase separation of the FUS low-complexity domain”. In: Nature Structural & Molecular Biology 26.7 (2019), pp. 637–648. doi: 10.1038/s41594-019-0250-x.

[59] Diana M. Mitrea et al. “Modulating biomolecular condensates: a novel approach to drug discovery”. In: Nature Reviews Drug Discovery 21.11 (2022), pp. 841–862. doi: 10.1038/s41573-022-00505-4.

[60] Gaurav Chauhan et al. “Crowder titrations enable the quantification of driving forces for macromolecular phase separation”. In: Biophysical Journal (2023), pp. 1–17. doi: 10.1016/j.bpj.2023.09.006.

[61] Ammon E Posey et al. “Biomolecular condensates are characterized by interphase electric potentials”. In: bioRxiv (2024), p. 2024.07.02.601783.

[62] Xue Wang et al. “Polyethylene Glycol Crowder’s Effect on Enzyme Aggregation, Thermal Stability, and Residual Catalytic Activity”. In: Langmuir 37.28 (2021), pp. 8474–8485. doi: 10.1021/acs.langmuir.1c00872.

[63] Christoph Klieber et al. “Mechanical spectra of glass-forming liquids. II. Gigahertz-frequency longitudinal and shear acoustic dynamics in glycerol and DC704 studied by time-domain Brillouin scattering”. In: Journal of Chemical Physics 138.12 (2013). doi: 10.1063/1.4789948.

[64] W. R. Moore. “Viscosity-Temperature Relationships for Dilute Solutions of High Polymers”. In: Nature 191.4795 (1961), pp. 1292–1293. doi: 10.1038/1911292b0.

[65] Han Yi Chou and Aleksei Aksimentiev. “Single-Protein Collapse Determines Phase Equilibria of a Biological Condensate”. In: Journal of Physical Chemistry Letters 11.12 (2020), pp. 4923–4929. doi: 10.1021/acs.jpclett.0c01222.

[66] Simon Alberti and Anthony A. Hyman. “Biomolecular condensates at the nexus of cellular stress, protein aggregation disease and ageing”. In: Nature Reviews Molecular Cell Biology 22.3 (2021), pp. 196–213. doi: 10.1038/s41580-020-00326-6.

[67] Adiran Garaizar et al. “Aging can transform single-component protein condensates into multiphase architectures”. In: Proceedings of the National Academy of Sciences 119.26 (2022), e2119800119. doi: 10.1073/pnas.2119800119.

[68] Maryam Alsadat Rad et al. “Micromechanical characterisation of 3D bioprinted neural cell models using Brillouin microspectroscopy”. In: Bioprinting 25 (2022), e00179. doi: 10.1016/j.bprint.2021.e00179.

[69] A. Bot, R. P. C. Schram, and G. H. Wegdam. “Brillouin light scattering from a biopolymer gel: hypersonic sound waves in gelatin”. In: Colloid & Polymer Science 273.3 (1995), pp. 252–256. doi: 10.1007/BF00657831.

[70] J. P. Jarry and G D Patterson. “Brillouin scattering from polyepoxide solutions and gels”. In: Macromolecules 14.5 (1981), pp. 1281–1284. doi: 10.1021/ma50006a026.

[71] Jennifer Illibauer et al. “Longitudinal viscosity of blood plasma for rapid COVID-19 prognostics”. In: medRxiv (2023). doi: 10.1101/2023.10.13.23297016.

[72] Clifford P. Brangwynne et al. “Germline P Granules Are Liquid Droplets That Localize by Controlled Dissolution/Condensation”. In: Science 324.5935 (2009), pp. 1729–1732. doi: 10.1126/science.1172046.

[73] Zakarya Benayad et al. “Simulation of FUS Protein Condensates with an Adapted Coarse-Grained Model”. In: Journal of Chemical Theory and Computation 17.1 (2021), pp. 525–537. doi: 10.1021/acs.jctc.0c01064.

[74] Mina Farag et al. “Condensates formed by prion-like low-complexity domains have small-world network structures and interfaces defined by expanded conformations”. In: Nature Communications 13.1 (2022), p. 7722. doi: 10.1038/s41467-022-35370-7.

[75] Giulio Tesei et al. “Accurate model of liquid–liquid phase behavior of intrinsically disordered proteins from optimization of single-chain properties”. In: Proceedings of the National Academy of Sciences 118.44 (2021). doi: 10.1073/pnas.2111696118.

[76] Moonseok Kim et al. “Shear Brillouin light scattering microscope”. In: Optics Express 24.1 (2016), p. 319. doi: 10.1364/OE.24.000319.

[77] Carlo Bevilacqua and Robert Prevedel. “Full-field Brillouin microscopy based on an imaging Fourier transform spectrometer”. In: (2024).

[78] Régis P. Lemaitre et al. “FlexiBAC: A versatile, open-source baculovirus vector system for protein expression, secretion, and proteolytic processing”. In: BMC Biotechnology 19.1 (2019), pp. 1–11. doi: 10.1186/s12896-019-0512-z.

[79] Raimund Schlüßler et al. “Mechanical Mapping of Spinal Cord Growth and Repair in Living Zebrafish Larvae by Brillouin Imaging”. In: Biophysical Journal 115.5 (2018), pp. 911–923. doi: 10.1016/j.bpj.2018.07.027.

[80] Raimund Schlüßler and others. BMicro. Version 0.8.1. 2023. url: https://github.com/BrillouinMicroscopy/BMicro.

[81] Giuseppe Antonacci et al. “Spectral broadening in Brillouin imaging”. In: Applied Physics Letters 103.22 (2013). doi: 10.1063/1.4836477.

[82] Matthäus Mittasch et al. “Non-invasive perturbations of intracellular flow reveal physical principles of cell organization”. In: Nature Cell Biology 20.3 (2018), pp. 344–351. doi: 10.1038/s41556-017-0032-9.

[83] Paul Müller. qpimage. Version 0.7.5. 2022. url: https://pypi.org/project/qpimage/0.7.5/.

[84] Timon Beck. DropletQPI. Version 0.1.0. 2023. url: https://github.com/GuckLab/DropletQPI.

[85] Paul Müller et al. “Accurate evaluation of size and refractive index for spherical objects in quantitative phase imaging”. In: Optics Express 26.8 (2018), p. 10729. doi: 10.1364/OE.26.010729.

[86] Paul Müller. qpsphere. Version 0.5.8. 2021. url: https://pypi.org/project/qpsphere/0.5.8/.

[87] Conrad Möckel et al. “Estimation of the mass density of biological matter from refractive index measurements”. In: bioRxiv (2023). doi: 10.1101/2023.12.05.569868.

[88] R. Barer. “Interference Microscopy and Mass Determination”. In: Nature 169.4296 (1952), pp. 366–367. doi: 10.1038/169366b0.

[89] H. G. Davies and M. H. F. Wilkins. “Interference Microscopy and Mass Determination”. In: Nature 169.4300 (1952), pp. 541–541. doi: 10.1038/169541a0.

[90] Huaying Zhao, Patrick H. Brown, and Peter Schuck. “On the distribution of protein refractive index increments”. In: Biophysical Journal 100.9 (2011), pp. 2309–2317. doi: 10.1016/j.bpj.2011.03.004.

[91] Daniel B. Allan et al. soft-matter/trackpy: Trackpy v0.5.0. Version v0.5.0. 2021.

[92] Simon Alberti et al. “A User’s Guide for Phase Separation Assays with Purified Proteins”. In: Journal of Molecular Biology 430.23 (2018). Phase Separation in Biology and Disease, pp. 4806–4820. doi: 10.1016/j.jmb.2018.06.038.

## References

[1] Lisa A. Romankiw and I Ming Chou. “Densities of aqueous sodium chloride, potassium chloride, magnesium chloride, and calcium chloride binary solutions in the concentration range 0. 5-6.1 m at 25, 30, 35, 40, and 45 °C”. In: Journal of Chemical & Engineering Data 28.3 (1983), pp. 300–305. doi: 10.1021/je00033a005.

[2] Arno Max Basedow, Klaus Heinrich Ebert, and Ursula Ruland. “Specific refractive index increments of dextran fractions of different molecular weights”. In: Die Makromolekulare Chemie 179.5 (1978), pp. 1351–1353. doi: 10.1002/macp.1978.021790523.

[3] Frank Doebert, Andreas Pfennig, and Matthias Stumpf. “Derivation of the Consistent Osmotic Virial Equation and Its Application to Aqueous Poly(ethylene glycol)-Dextran Two-Phase Systems”. In: Macromolecules 28.23 (1995), pp. 7860–7868. doi: 10.1021/ma00127a037.

[4] Conrad Möckel et al. “Estimation of the mass density of biological matter from refractive index measurements”. In: bioRxiv (2023). doi: 10.1101/2023.12.05.569868.

[5] Robert Prevedel et al. “Brillouin microscopy: an emerging tool for mechanobiology”. In: Nature Methods 16.10 (2019), pp. 969–977. doi: 10.1038/s41592-019-0543-3.

[6] Raimund Schlüßler et al. “Mechanical Mapping of Spinal Cord Growth and Repair in Living Zebrafish Larvae by Brillouin Imaging”. In: Biophysical Journal 115.5 (2018), pp. 911–923. doi: 10.1016/j.bpj.2018.07.027.

[7] Giuseppe Antonacci et al. “Spectral broadening in Brillouin imaging”. In: Applied Physics Letters 103.22 (2013). doi: 10.1063/1.4836477.

[8] Maurizio Mattarelli, Massimo Vassalli, and Silvia Caponi. “Relevant Length Scales in Brillouin Imaging of Biomaterials: The Interplay between Phonons Propagation and Light Focalization”. In: ACS Photonics (2020). doi: 10.1021/acsphotonics.0c00801.

[9] Jennifer Illibauer et al. “Longitudinal viscosity of blood plasma for rapid COVID-19 prognostics”. In: medRxiv (2023). doi: 10.1101/2023.10.13.23297016.

[10] Raimund Schüßler et al. “Correlative all-optical quantification of mass density and mechanics of subcellular compartments with fluorescence specificity”. In: eLife 11 (2022). Ed. by Rohit V Pappu and Anna Akhmanova, e68490. doi: 10.7554/eLife.68490.

[11] Huaying Zhao, Patrick H. Brown, and Peter Schuck. “On the distribution of protein refractive index increments”. In: Biophysical Journal 100.9 (2011), pp. 2309–2317. doi: 10.1016/j.bpj.2011.03.004.

[12] Gertrude E. Perlmann and L. G. Longsworth. “The Specific Refractive Increment of Some Purified Proteins”. In: Journal of the American Chemical Society 70.8 (1948), pp. 2719–2724. doi: 10.1021/ja01188a027.

[13] Tsutomu Arakawa. “Calculation of the Partial Specific Volumes of Proteins in Concentrated Salt, Sugar, and Amino Acid Solutions”. In: The Journal of Biochemistry 100.6 (1986), pp. 1471–1475. doi: 10.1093/oxfordjournals.jbchem.a121853.

[14] Ingrid Pilz and Gertrud Czerwenka. “The partial specific volume of various proteins and its dependence on concentration and temperature of the solution”. In: Die Makromolekulare Chemie 170.1 (1973), pp. 185–190. doi: 10.1002/macp.1973.021700115.

